# A systematic review of the diversity and virulence correlates of metastrongyle lungworms in marine mammals

**DOI:** 10.1101/2023.02.03.527020

**Authors:** Jared R. Fischbach, Mauricio Seguel

## Abstract

Metastrongyle lungworms could be particularly detrimental for diving animals such as marine mammals, however little is known of the drivers of pathogenic host-parasite relationships in this group. This systematic review analyzed the diversity of metastrongyles in marine mammals and the host and parasite traits associated with virulence. There have been at least 40 species of metastrongyles described in 66 species of marine mammals. After penalization for study biases, *Halocercus hyperoodoni*, *Otostrongylus circumlitus*, *Parafilaroides gymnurus*, *Halocercus brasiliensis*, and *Stenurus minor* were the metastrongyles with the widest host range. Most studies (80.12%, n=133/166) reported that metastrongyles caused bronchopneumonia, while in the cardiovascular system metastrongyles caused vasculitis in nearly half of the studies (45.45%, n=5/11) that assessed these tissues. Metastrongyles were associated with otitis in 23.08% (n=6/26) of the studies. Metastrongyle infection was considered a potential contributory to mortality in 44.78% (n=90/201) of the studies while 10.45% (n=21/201) of these studies considered metastrongyles the main cause of death. Metastrongyle species with a wider host range were more likely to induce pathogenic effects. Metastrongyles can cause significant tissue damage and mortality in marine mammals although virulent host-parasite relationships are dominated by a few metastrongyle species with wider host ranges.

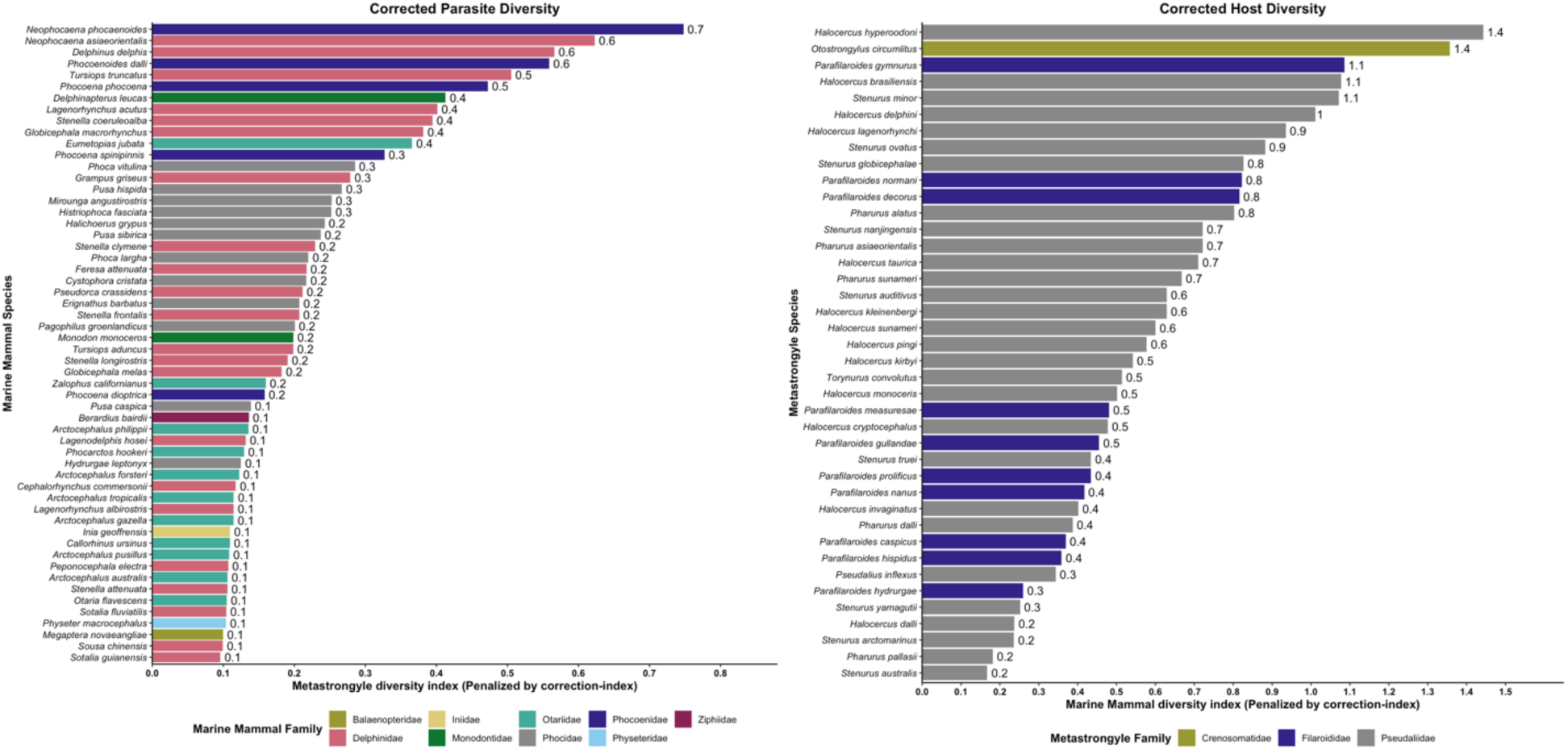

## 1. Introduction

Metastrongyloidea (Nematoda: Strongylida), also known as lungworms or metastrongyles, are common parasites of the respiratory, cardiovascular, and rarely the auditory system of mammals (Measures, 2001). The definitive hosts become infected through ingestion of larval stages within invertebrate intermediate hosts or free on pasture, therefore, metastrongyle infection can be quite common among free-foraging domestic and wild mammals (Measures, 2001; Regassa *et al*., 2010; Niebuhr *et al*., 2021). Like other helminth infections, the pathogenic effects of metastrongyloidiasis can be broad with impacts that range from subclinical infection to bronchopneumonia and death (Measures, 2001; Forbes, 2021). The reasons for these variable outcomes are not fully understood but site of infection (e.g. upper vs lower airways), burden and host immune status have been indicated as drivers of pathogenic outcomes (Forbes, 2021). However, it is unclear if certain host-parasite relationships are more detrimental for the host and if these more virulent associations obey to ecological factors that could illuminate conserved drivers of pathogenicity in these nematodes. Interestingly, the virulence of metastrongyles could be higher for animal species that require optimal respiratory and auditory function for fitness, such as marine mammals (Bergeron *et al*., 1997b; Measures, 2001; Seibel *et al*., 2010). Marine mammals rely on cardiovascular and lung capacity for foraging dives and the otic apparatus of cetaceans is critical for communication and echolocation in the water column (Bergeron *et al*., 1997b; Measures, 2001; Seibel *et al*., 2010). Therefore, metastrongyles parasitism of these host compartments could be associated with disease, stranding and death (Bergeron *et al*., 1997b; Measures, 2001; Seibel *et al*., 2010), making this animal group an interesting model to test conserved correlates of virulence among metastrongyle species. Nevertheless, metastrongyles are one of the most diagnosed parasites in marine mammals, and in many circumstances are considered incidental findings of little to no pathologic significance (Geraci and St. Aubin, 1987). This conflicting evidence could be related to negative metastrongyle effects limited to specific host-parasite relationships, parasite traits, environmental conditions, or the research methods employed to assess the health impacts. However, a systematic assessment of the factors that could modulate the health impacts of metastrongyles has not been reported.

Three families of metastrongyles are prominently found in marine mammals, spanning seven genera, with a considerable diversity of species within each genus. Despite this diversity, metastrongyles share similar life histories and host exploitation strategies in marine mammals. Most aquatic metastrongyle species require an obligate intermediate host, usually an invertebrate, where several larval stages develop until reaching an infective stage (Dailey, 1970a, 2006a). These infective larvae are consumed by the definitive host and then migrate from the gastrointestinal system to the respiratory or cardiovascular system where sexual maturity and reproduction occurs, releasing embryonated-larvated eggs or larvae (Taylor *et al*., 2015). Additionally, in some species, maternal lungworms infect neonates through the placenta and/or milk (Dailey *et al*., 1991a). In the airways and respiratory sinuses or cardiovascular system, metastrongyles feed on epithelial-endothelial cells, host debris and occasionally blood (Dailey, 1970a; Fauquier *et al*., 2010; Rhyan *et al*., 2018a). These processes of migration through tissues, feeding and releasing eggs and larvae illicit variable immune responses in the host (Baylis and Daubney, 1925; Fauquier *et al*., 2010; Groch *et al*., 2020a). This immune response is usually directed against parasite cuticle, larvae, eggs or parasite remains and usually consist in the attraction of macrophages, eosinophils and lymphocytes to the site of infection (Fauquier *et al*., 2010; Zafra *et al*., 2015a; Seguel *et al*., 2020a; Groch *et al*., 2020a). The degree of this inflammatory response and its impact on health status can be quite variable ranging from minimal to severe bronchopneumonia (Fauquier *et al*., 2010; Seguel *et al*., 2020a; Groch *et al*., 2020a), being the drivers of the variable responses poorly understood.

There is conflicting evidence regarding the connection between metastrongyle infections and cetacean strandings (Arbelo *et al*., 2013). Currently, there is no consensus on whether inflammation in the otic apparatus, elicited by a presence of metastrongyle parasites, can impair hearing or echolocation to a significant enough degree to be considered a contributory factor in cetacean strandings (Geraci and St. Aubin, 1987; Wohlsein *et al*., 2019a). Similarly, the presence and feeding behavior of the nematodes in bronchi and lung parenchyma facilitates secondary bacterial infections with conflicting documented outcomes, ranging from little importance for lung physiology to fatal bronchopneumonia, especially when parasites are found in the lower respiratory system (Siebert *et al*., 2001; Lehnert *et al*., 2010; Ulrich *et al*., 2015, 2016; Reckendorf *et al*., 2021). This conflicting evidence could be related to several host and parasite traits, although across biological systems, parasite biomass, usually assessed in terms of burden, is a common driver of negative health outcomes (Pedersen and Fenton, 2015; Seguel and Gottdenker, 2017). Similarly, parasite and host sizes play an important role in assessing total parasite biomass and can sometimes serve as useful proxies of biomass to identify particularly detrimental host-parasite relationships across studies (Seguel and Gottdenker, 2017). Another potential factor impacting health outcomes across host-parasite systems is the capacity of certain helminths to parasitize multiple host species. Parasites with a wider host range can potentially become more virulent because decreases in host fitness due to infection can be compensated by infecting other susceptible species (Poulin, 2011; Schmid-Hempel, 2021). However, the role of these host and parasite traits on the health impact of metastrongyles is not well defined.

The objectives of this systematic review were to i) summarize existing literature concerning relationships between marine mammals and metastrongyle parasites, ii) assess the relative host and parasite diversity of infections iii) outline the pathology and health consequences that result from metastrongyle infections in marine mammals, with assessment of the potential drivers of these effects.

## 2. Materials and Methods

### 2.1 Searching methods and inclusion criteria

A systematic literature review on metastrongyle parasites in marine mammals was conducted on July 6, 2021. The search was conducted using Google Scholar and PubMed, following the best practices outlined by Haddaway and Watson (2016) to conduct an aggregative systematic review in the field of parasitology (Haddaway and Watson, 2016) and the recommendations outlined in the “Preferred Reporting Items for Systematic Reviews and Meta-Analyses” (Page *et al*., 2021).

The search terms used for the literature search on PubMed were as follows: “((Marine) OR (Cetacean) OR (Delphinidae) OR (Phocidae) OR (Phocoenidae) OR (Otariidae) OR (Monodontidae) OR (Mustelidae) OR (Trichechidae) OR (Odobenidae) OR (Ursidae) OR (Odobenidae) OR (Balaenidae) OR (Neobalaenidae) OR (Eschrichtiidae) OR (Balaenopteridae) OR (Physeteridae) OR (Kogiidae) OR (Ziphiidae) OR (Platanistidae) OR (Iniidae) OR (Lipotidae) OR (Pontoporiidae) OR (Dugongidae) OR (Seal) OR (Porpoise) OR (Otter) OR (Dolphin) OR (Sea lion) OR (Whale) OR (Walrus) OR (Manatee)) AND ((Metastrongyle) OR (Metastrongylus) OR (Metastrongyloidea) OR (Metastrongylidae) OR (Parafilaroides) OR (Stenurus) OR (Halocercus) OR (Pseudaliidae) OR (Filaroididae) OR (Crenosomatidae) OR (Otostrongylus) OR (Skrjabingylus))”.

The search terms used for the literature search on Google Scholar were reduced due to limitations in the advanced search software, notably a 256-character limit on searches. The search terms used for Google Scholar were as follows: “((Metastrongyle) OR (Metastrongylus) OR (Metastrongyloidea) OR (Metastrongylidae) OR (Parafilaroides) OR (Stenurus) OR (Halocercus) OR (Pseudaliidae) OR (Filaroididae) OR (Crenosomatidae) OR (Otostrongylus) OR (Skrjabingylus)) AND ((Marine) OR (Cetacean))”.

The search on PubMed yielded 120 results and the search on Google Scholar yielded 2110 results, excluding citations. As Google Scholar can only display 1000 results per search, only the 1000 articles deemed most relevant by the software were manually reviewed for inclusion in the study, in addition to the 120 articles found via PubMed. To ensure all recently published literature was consulted, a secondary search on google scholar was conducted with the same search terms, filtering to only include articles published from 2017 and onward. This secondary search yielded 380 results, excluding citations, which were also manually reviewed for inclusion in the study. All conference archives (e.g. IAAAM), theses not published in peer-reviewed journals, and posters were excluded from this study. The full texts for the articles isolated through literature search results were obtained and manually reviewed by the two authors, being included in the study if they met the following inclusion criteria: i) The host species was any wild marine mammal, with captive wild animals only included if the transmission of parasites was stated to have occurred naturally, prior to their capture. ii) The parasite species belonged to a genus within the metastrongyle superfamily, including studies that described the presence of parasitic eggs and larvae provided they could be positively identified to at least the genus level (studies that described parasites as “lungworms” or “nematodes” without specificity regarding genera were excluded). iii) The article serves as the primary source describing a relationship between a marine mammal host and metastrongyle parasite. Duplicate articles, existing in both the Google Scholar and PubMed searches were manually removed. All non-primary sources which were found through the initial search terms were screened, and citations were followed, leading to the inclusion of 15 additional articles which met the inclusion criteria. Later, the World Register of Marine Species (WoRMS, https://www.marinespecies.org), The Society for Marine Mammalogy (https://marinemammalscience.org), and the Natural History Museum Host-Parasite database (https://www.nhm.ac.uk/research-curation/scientific-resources/taxonomy-systematics/host-parasites/) were consulted to ensure all known unique relationships were reported in this study (WoRMS Editorial Board, 2022). A total of 35 articles could not be acquired in their full text form for inclusion in data analysis, but the relationships they report are known through prior citations in peer-reviewed journals or available abstracts. The articles which met the inclusion criteria were further evaluated by the two authors, with significant attributes documented on a master spreadsheet. Many included articles, including the articles not available in full-text form, report multiple different host-parasite relationships, thus independent rows were created on the master spreadsheet for each described host-parasite relationship (n=481), allowing for relationship-specific attributes to be evaluated. The accepted common names and scientific names provided for host species in this study are taken from the Society for Marine Mammalogy Committee on Taxonomy (WoRMS Editorial Board, 2022).

A total of 207 full text publications met the inclusion criteria for this study, with 114 appearing only on Google Scholar, 6 appearing only on PubMed, 72 appearing on both Google Scholar and PubMed, and 15 coming from other sources. Data was collected on all included full-text publications, including whether they assessed pathogenic effect. An article was deemed to assess pathogenic effect if it described some biological impact associated with a presence of metastrongyle parasites in the host, or if it explicitly stated no pathogenic effect was associated with the parasitism. Further data was collected on whether full-text publications which met the inclusion criteria mentioned strandings or mortality. For the collection of this data, an article was deemed to mention strandings if it contained the following terms (not including in the citations): “stranded” or “stranding” or “strandings”. For an article to have been deemed to mention mortality, it must have contained the following terms (not including in the citations): “mortality” or “mortalities”. The inclusion terms for determining which articles mention stranding or mortality were isolated through word searching in adobe acrobat, with technologically non-conforming articles screened manually. Later, the studies mentioning mortality or stranding were assessed to ensure they discussed these effects in the context of the study’s findings. The parasite collection methodology was also noted for each included full-text publication. Within the articles which met the inclusion criteria, the geographical location(s) were recorded for 437 documented relationships and the anatomical location(s) of the parasites were recorded in 371 documented relationships. We further calculated the geographical dispersion of the 102 unique relationships between marine mammals and metastrongyles.

For each documented relationship, data was also recorded on any reported pathogenic effects (n=201). Pathogenic effects can be described as some biological impact on the host associated with the presence of metastrongyle parasites. In cases where it was explicitly stated that no pathogenic effect was associated with parasitism, this was recorded as well. Of the 101 articles that assessed pathogenic effect, there were a total of 201 relationships in which attributes describing a pathogenic effect or lack thereof were documented.

Between the 207 full-text publications which met our inclusion criteria, and the additional 35 publications we could not retrieve in full-text form, a total of 481 host-parasite relationships are documented. For each of these relationships, data was collected on their respective species taxonomy. Next, the frequency at which each metastrongyle parasite species appears in the literature, in the context of relationships with marine mammal hosts was recorded. For the purposes of frequency calculations, parasites described only to the genus level were excluded. The frequency at which marine mammal hosts appear in the literature, in the context of relationships with metastrongyle parasites was recorded in the same manner. For frequency calculations, hosts described only to the genus level were excluded. For the purposes of all diversity calculations, only relationships where both the host and parasite were described to the species level were considered. Finally, host diversity was calculated as a measure of the diversity of host species a metastrongyle species can infect, and parasite diversity was calculated as the diversity of metastrongyle species a marine mammal species can host.

To provide a metric by which the size of a host could be used as a predictor of pathogenic effects, host body mass was calculated by dividing the median weight in kilograms by the median length in centimetres for each host species. Median host measurements were compiled as the median of both genders from a variety of sources based on availability in the following order of priority: 1) World Register of Marine Species (WoRMS, https://www.marinespecies.org); 2) Society for Marine Mammalogy (https://marinemammalscience.org/science-and-publications/species-information/facts/); 3) National Oceanic and Atmospheric Administration (NOAA, https://www.fisheries.noaa.gov/species-directory); 4) Animal Diversity Web (ADW, https://animaldiversity.org). In cases where data on host size was not available through those four sources, the median measurements of hosts documented during gross necropsy in articles which met the inclusion criteria of this study were used.

Finally, we produced an up-to-date checklist of all known relationships concerning metastrongyles in marine mammals, which is included in table 1, complete with information on anatomical ranges documented for each relationship. The relationships documented are found within the 207 articles included in this study or within the 35 articles that could not be acquired in their full text form for inclusion in data analysis, provided the relationships they report are known through prior citations in peer-reviewed journals.

**Table 1:**
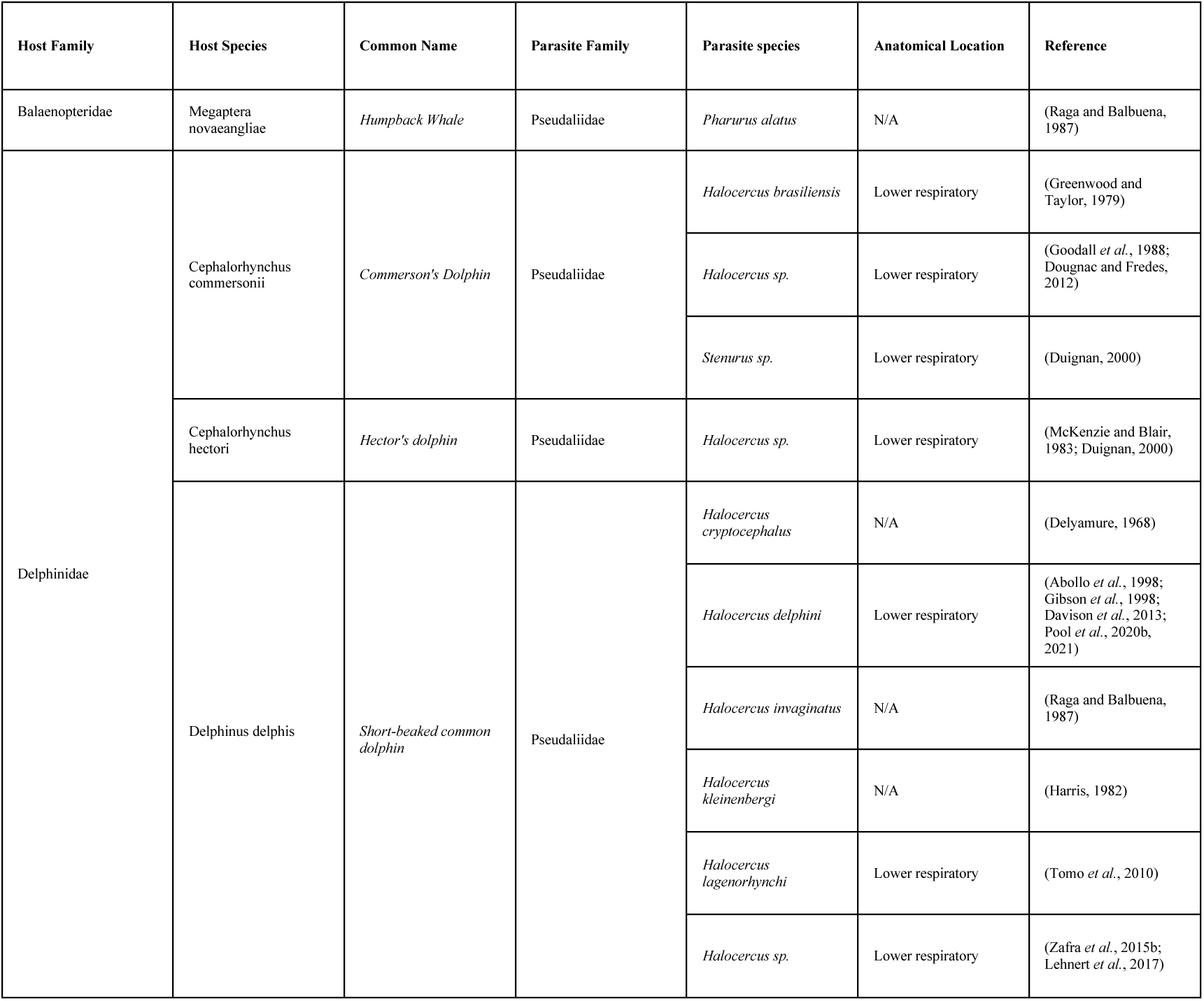
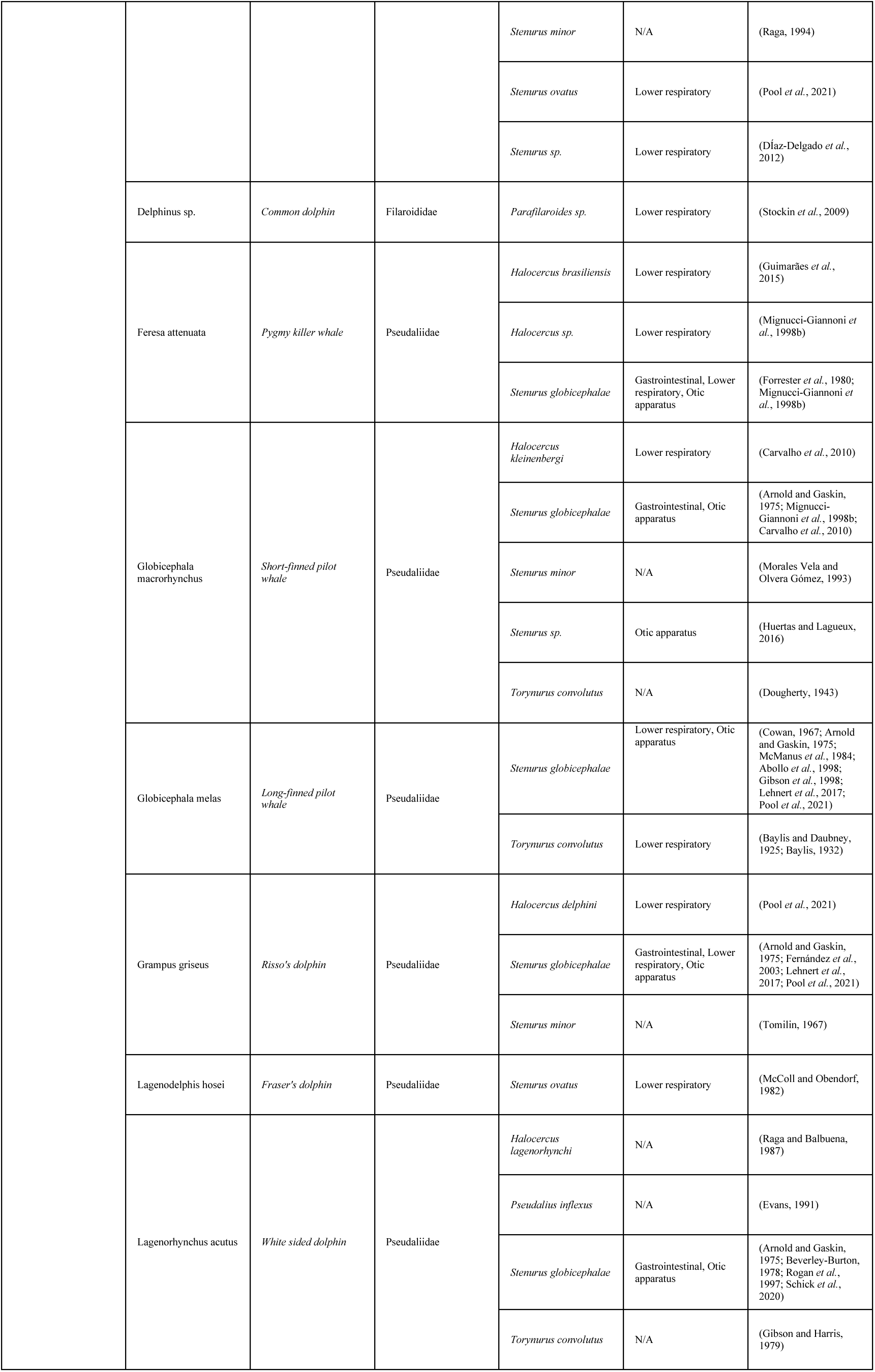
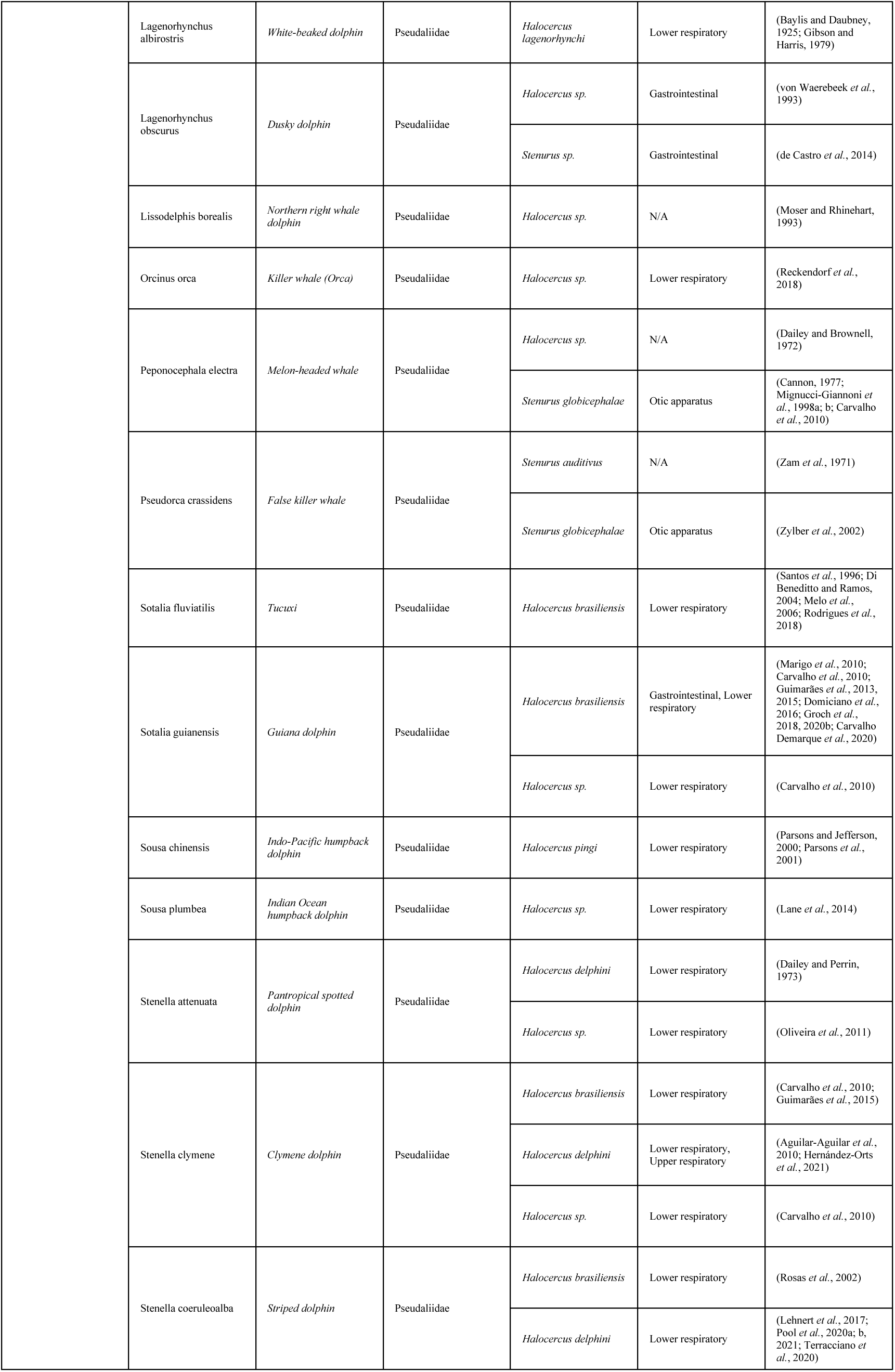
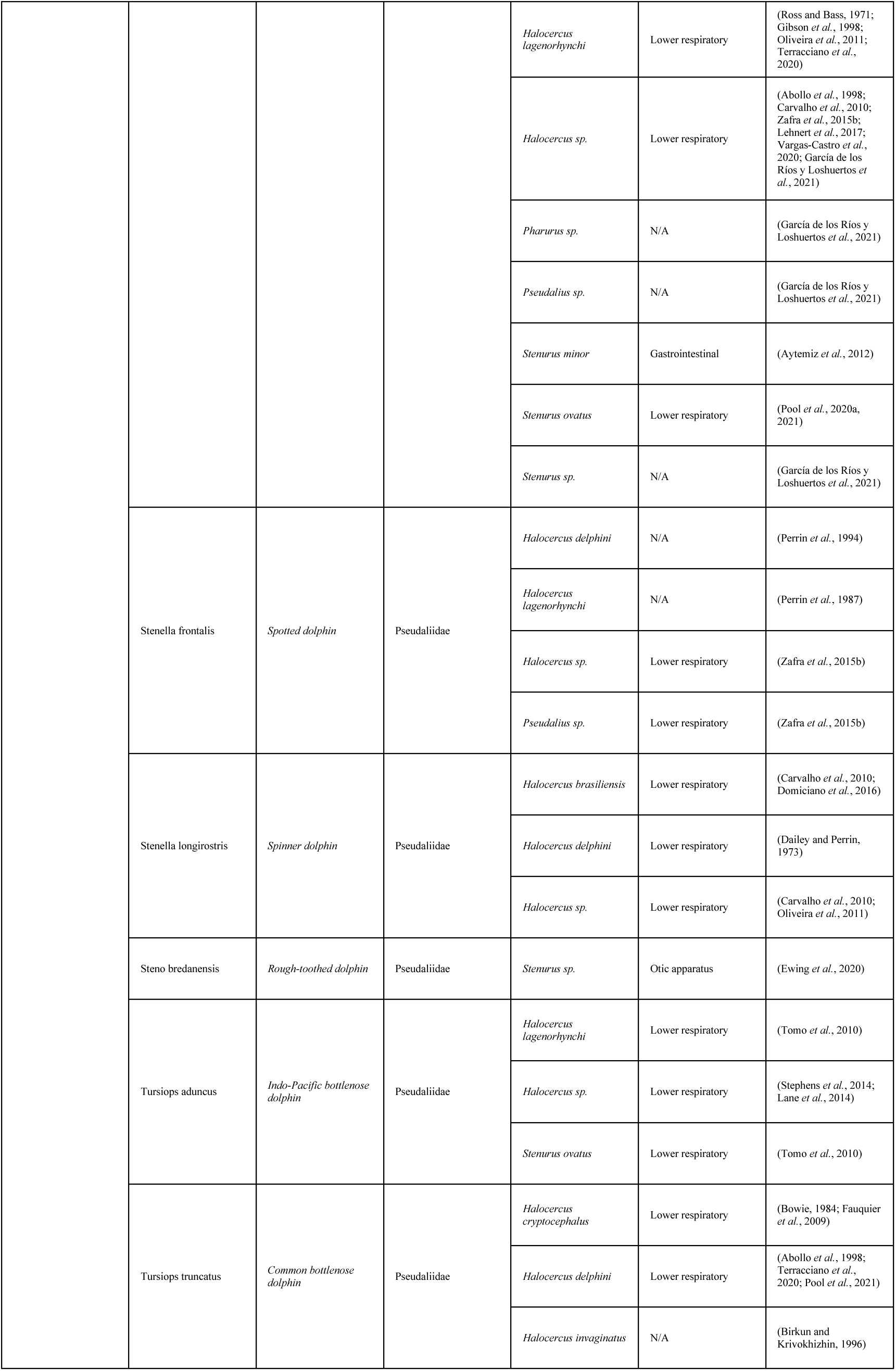
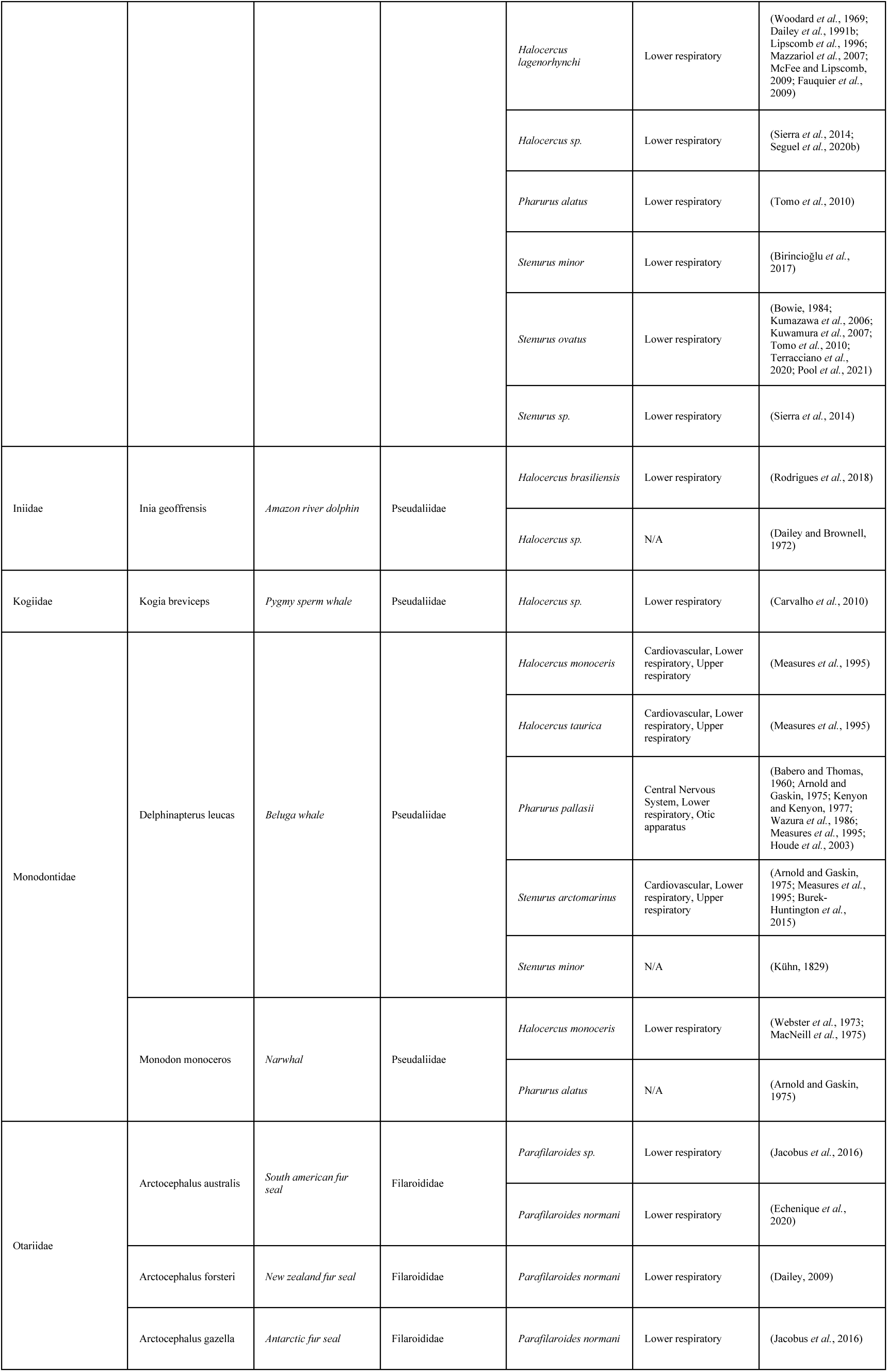
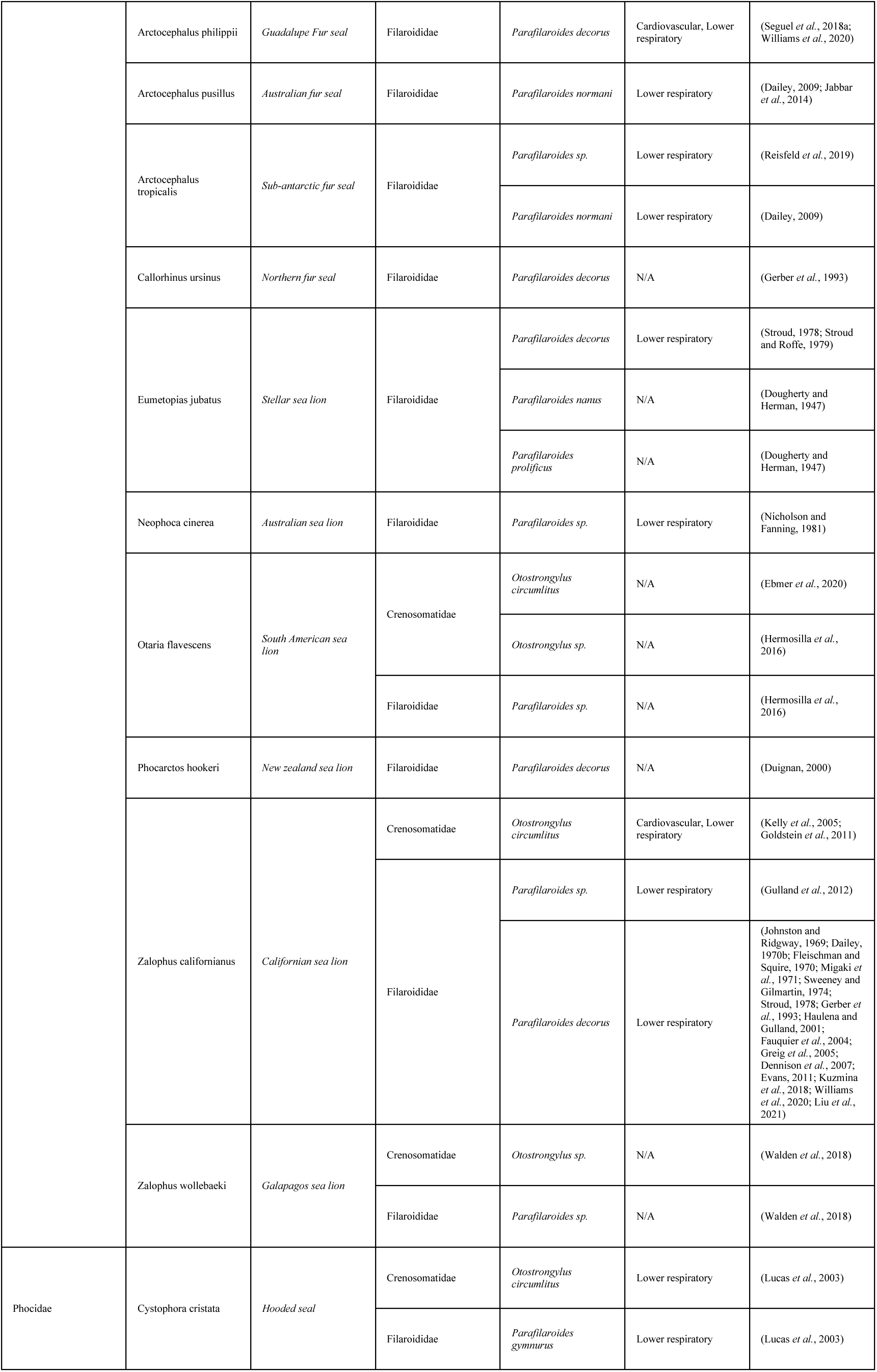
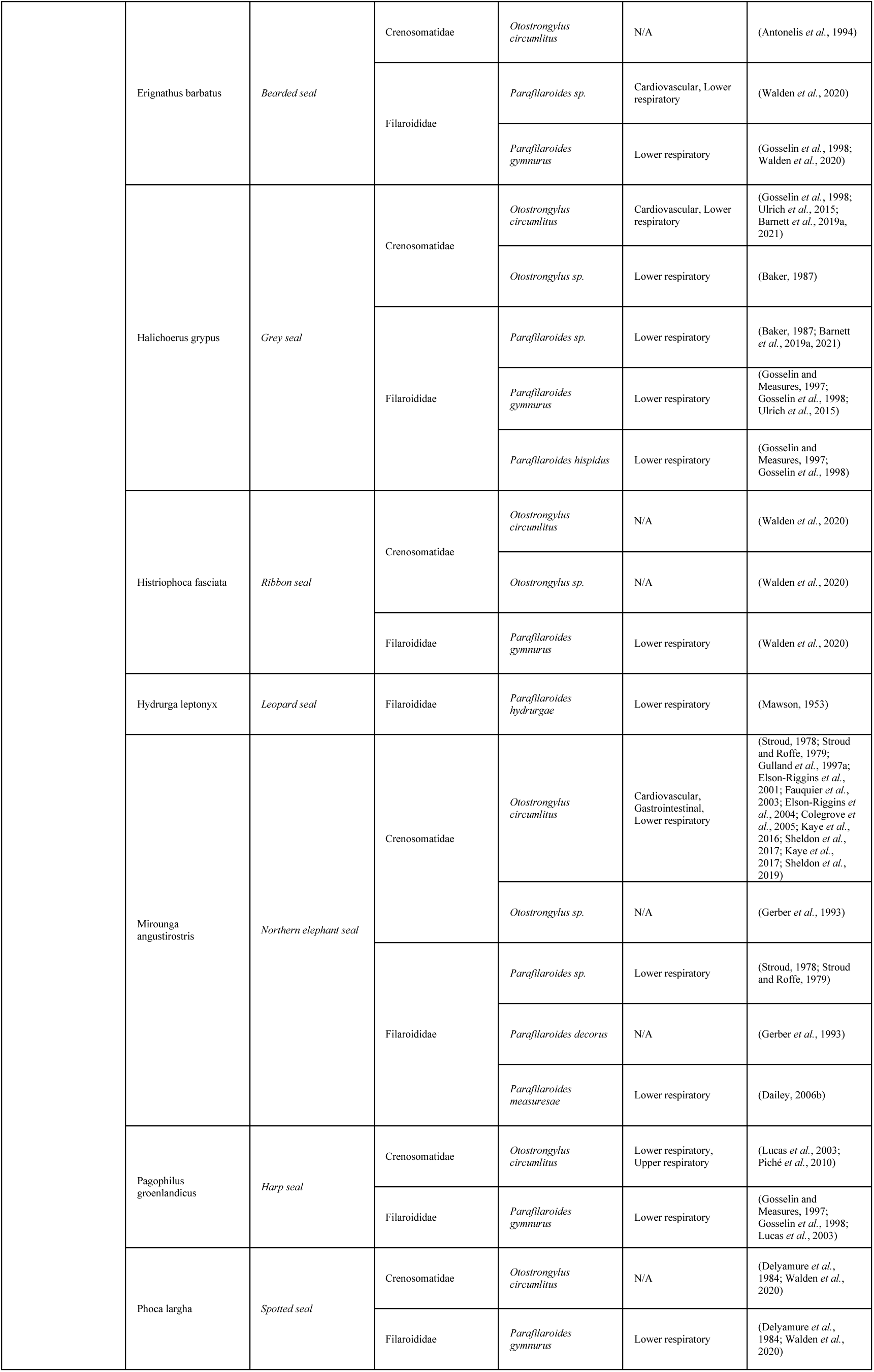
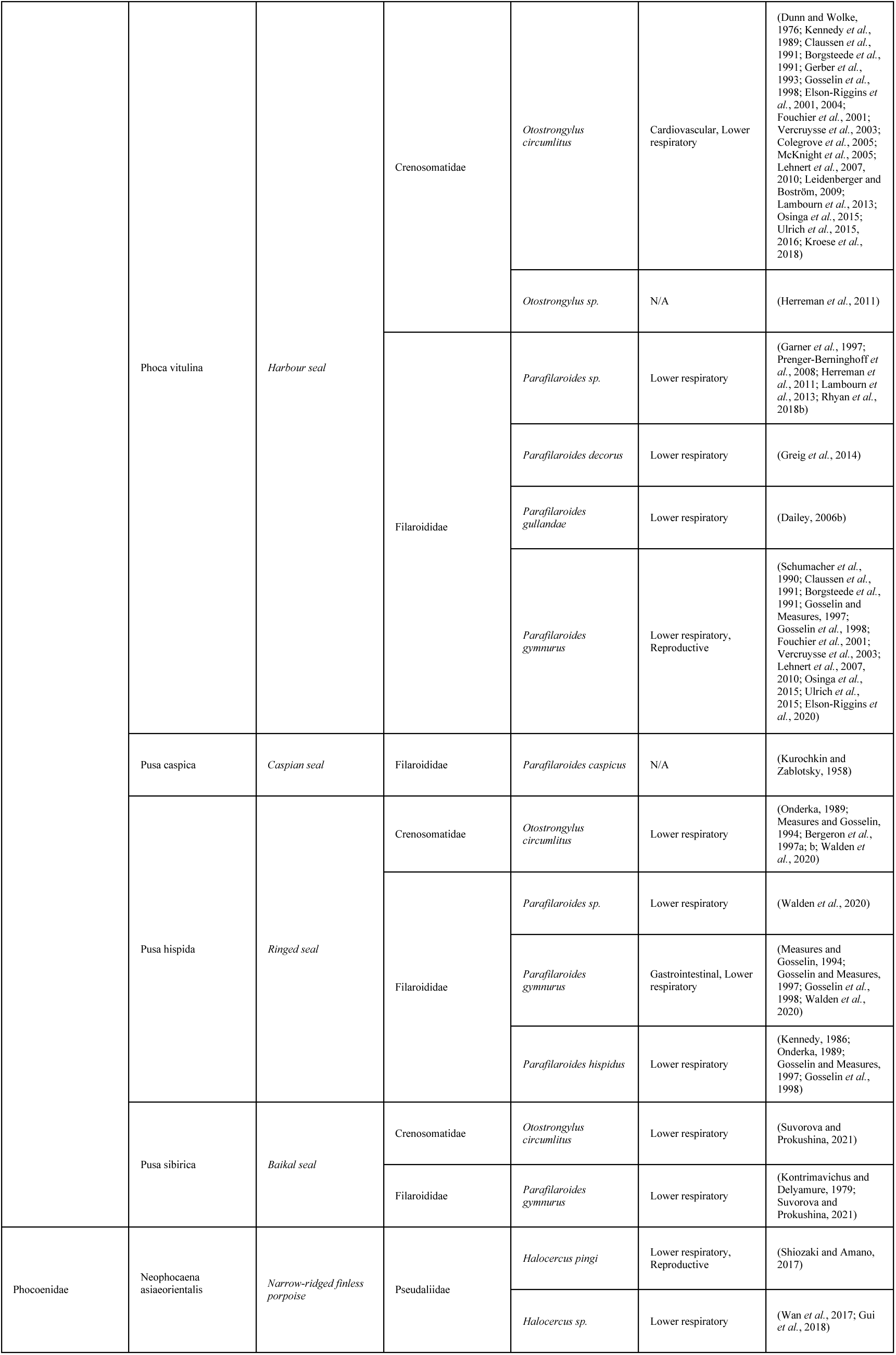
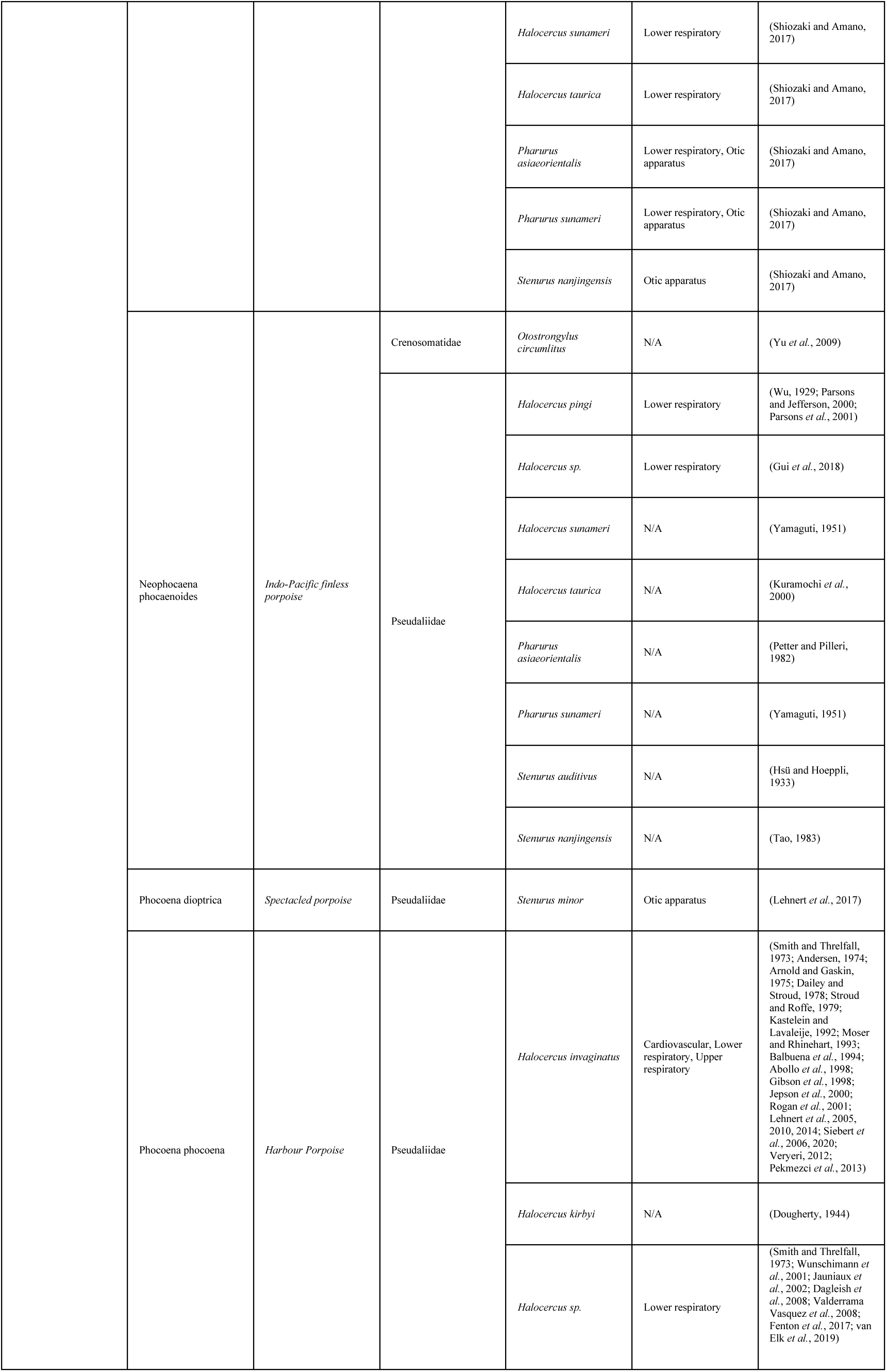
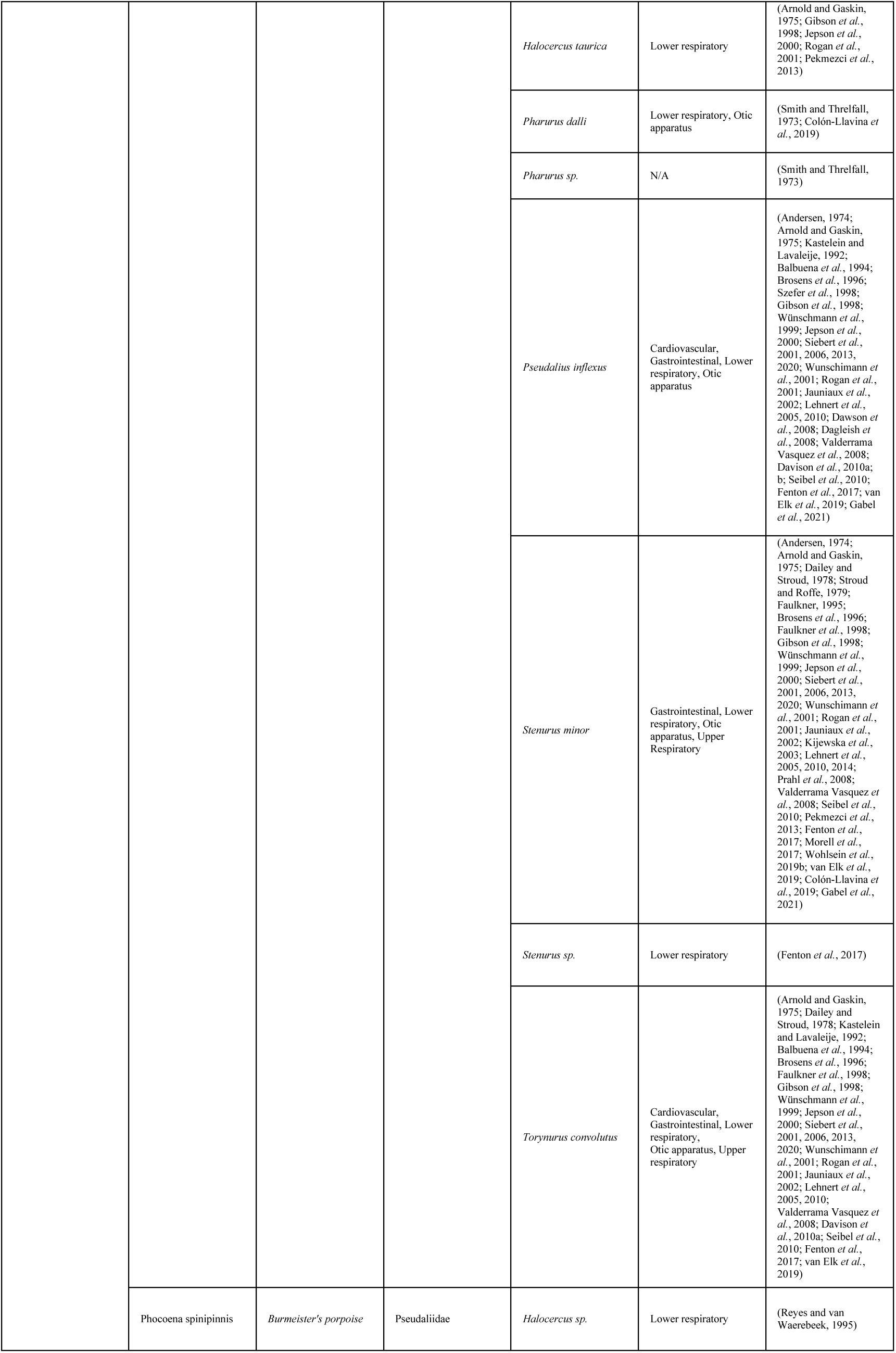
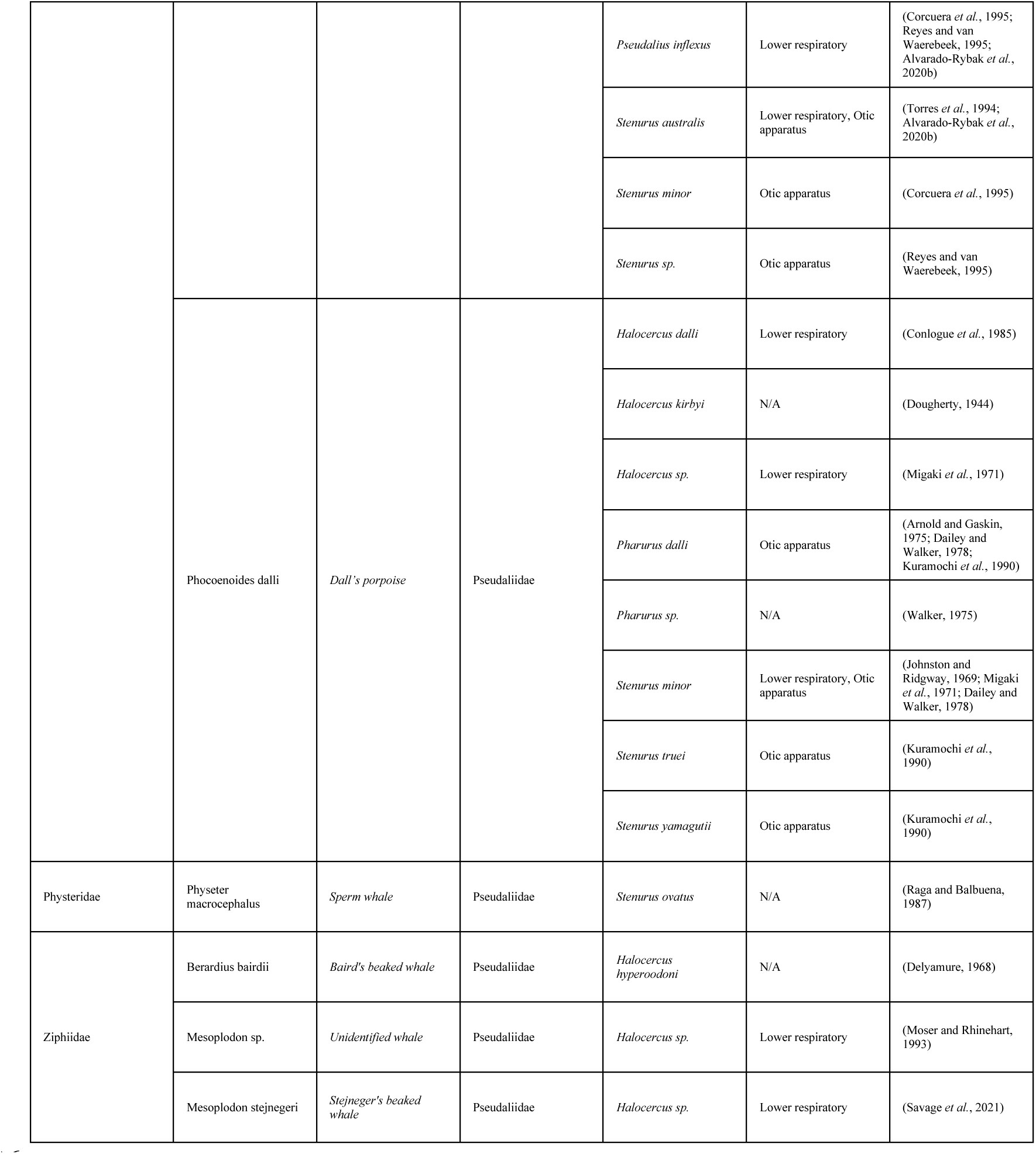
Checklist of all known relationships concerning metastrongyles in marine mammals, complete with information on anatomical ranges documented for each relationship.

### 2.2 Data management and analyses

When assessing the relative host and parasite diversities, corrections were made for literature and intra-study biases which arose from non-uniform sampling efforts. We used a modified approach, based previously published methods (Nunn *et al*., 2003; Ezenwa *et al*., 2006; Seguel and Gottdenker, 2017), to create a bias penalization index for each host and parasite species. To do this, we searched scientific names of each host and parasite species individually on Google Scholar and the number of hits was recorded. The number of hits for each species was then multiplied by the number of studies that both met our inclusion criteria and contained mentions of a relationship concerning that respective species. The natural logarithm of these products was then taken, and the result constitutes our penalization index for each species study bias (correction-index). The resulting corrected diversity indexes were plotted using frequency plots constructed in the R package “ggplo2”.

We explored the predictors of pathogenic effects among different host parasite interactions using binomial generalized linear mixed effects models (GLMMs) with the restricted maximum likelihood fitted using the R package “glmmTMB” (Brooks *et al*., 2017). Although this method can estimate positively biased odds ratios for binary outcomes in metadata, these are usually more problematic for smaller samples sizes and alternative methods for the analysis of binary metadata present similar issues (Stijnen *et al*., 2010; Bakbergenuly and Kulinskaya, 2018). We used corrected host diversity, corrected parasite diversity, and host body size as predictors for the presence or absence of pathogenic effects. Study ID was added as a random effect to account for studies documenting multiple host-parasite relationships with their respective outcomes. For each documented host-parasite relationship, with both the host and parasite specified to the species level and for which pathogenic effect was assessed (n=165), binomial GLMMs for the presence or absence of pathogenic effect were fit. A second model was fit to exclusively assess the presence or absence of respiratory specific effect (bronchopneumonia or bronchitis). For the respiratory specific effect model, each documented host-parasite relationship was included if the host and parasite were specified to the species level, pathogenic effect was assessed, and the presence of the parasite was documented in either the upper or lower respiratory system (n=134). We attempted to fit cardiovascular specific effect (DIC, thrombosis, vasculitis, hemorrhage, endocarditis) (n=11) and otic apparatus specific effect (sinusitis, otitis, otic metaplasia) (n=24) models using the same method as for respiratory specific effect but were unable because the small sample size affected model convergence. In both the full and respiratory specific models, we included interaction effects between the corrected host diversity, corrected parasite diversity, and body size predictors terms to account for individual permissiveness of host species to parasite species as well as individual pervasiveness of parasite species to host species.

### 2.3 Discrepancies in the literature

Hermosilla, C., et al. (2015) reports *Stenurus sp*. in sei whale (*Balaenoptera borealis*), blue whale (*Balaenoptera musculus*), and fin whale (*Balaenoptera physalus*) as non-primary findings, however we could not identify a primary source of these relationships (Hermosilla *et al*., 2015). Lehnert, K., et al. (2019) reports *Pharurus alatus* as a parasite of short-beaked common dolphin (*Delphinus delphis*) citing Tomo, I., et al. (2010) (Tomo *et al*., 2010; Lehnert *et al*., 2019). Upon reviewing Tomo, I., et al. (2010), it does not appear to include this relationship (Tomo *et al*., 2010). The World Register of Marine Species (WoRMS) documents *Halocercus taurica* in of short-beaked common dolphin (*Delphinus delphis*) and *Halocercus hyperoodoni* in northern bottlenose whale (*Hyperoodon ampullatus*), however primary sources could not be identified for either of these relationships (WoRMS Editorial Board, 2022). Measures, L.N. (2001) documents *Halocercus pingi* in Dall’s porpoise (*Phocoenoides dalli*) as a non-primary finding, however we could not find a primary source for this relationship (Measures, 2001). Three (n=3) additional publications document relationships that were not specified to the genus level (Giorda *et al*., 2017; Reckendorf *et al*., 2021; van Wijngaarden *et al*., 2021).

## 3. Results

At least 40 species of metastrongyles have been described in 66 marine mammal host species. Most metastrongyle species parasitize multiple host species, with a total of 213 unique host-metastrongyle relationships recorded (table 1).

### 3.1 Characterization of studies

The mean publication year of the 207 articles which met the inclusion criteria was 2003, with the distribution of publication years skewed towards the most recent decade (Figure 1). For each of the articles which met the inclusion criteria, 48.88% (n=101/207) assessed pathogenic effect. Additionally, 78.74% (n=163/207) of studies mentioned stranding and 43.48% (n=90/207) of articles mentioned mortality as potentially related to lungworm infection. Gross necropsy is the predominant parasite collection method for metastrongyles in marine mammals with 92.27% (n=191/207) of the studies using this as the only method of collection. The next most common method of collection was, fecal sampling, in 6.28% (n=13/207) of the studies (Figure 2).

**Figure 1:**
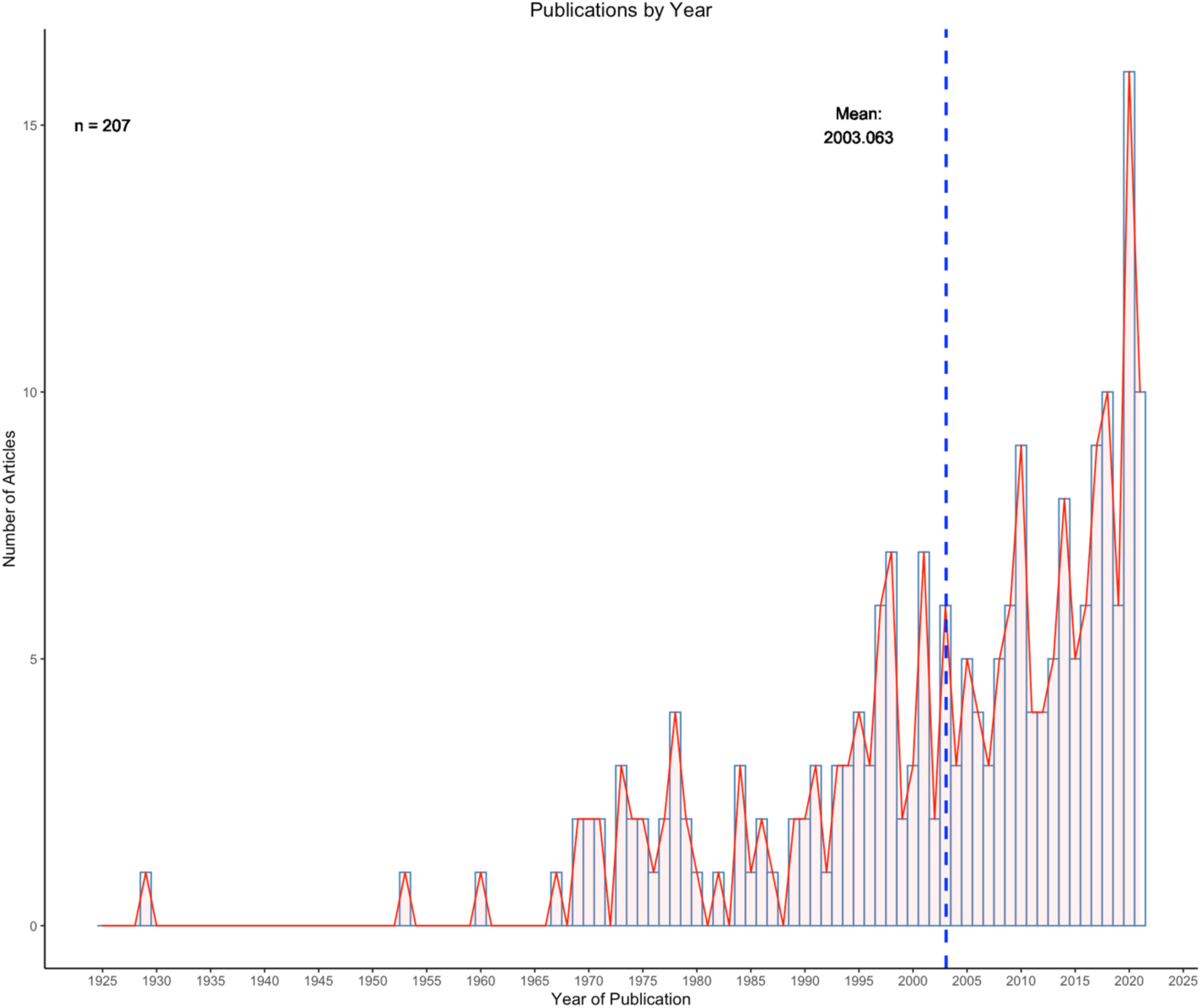
Distribution of publication year for articles which met the inclusion criteria of this study. The mean year of publication was 2003.

**Figure 2:**
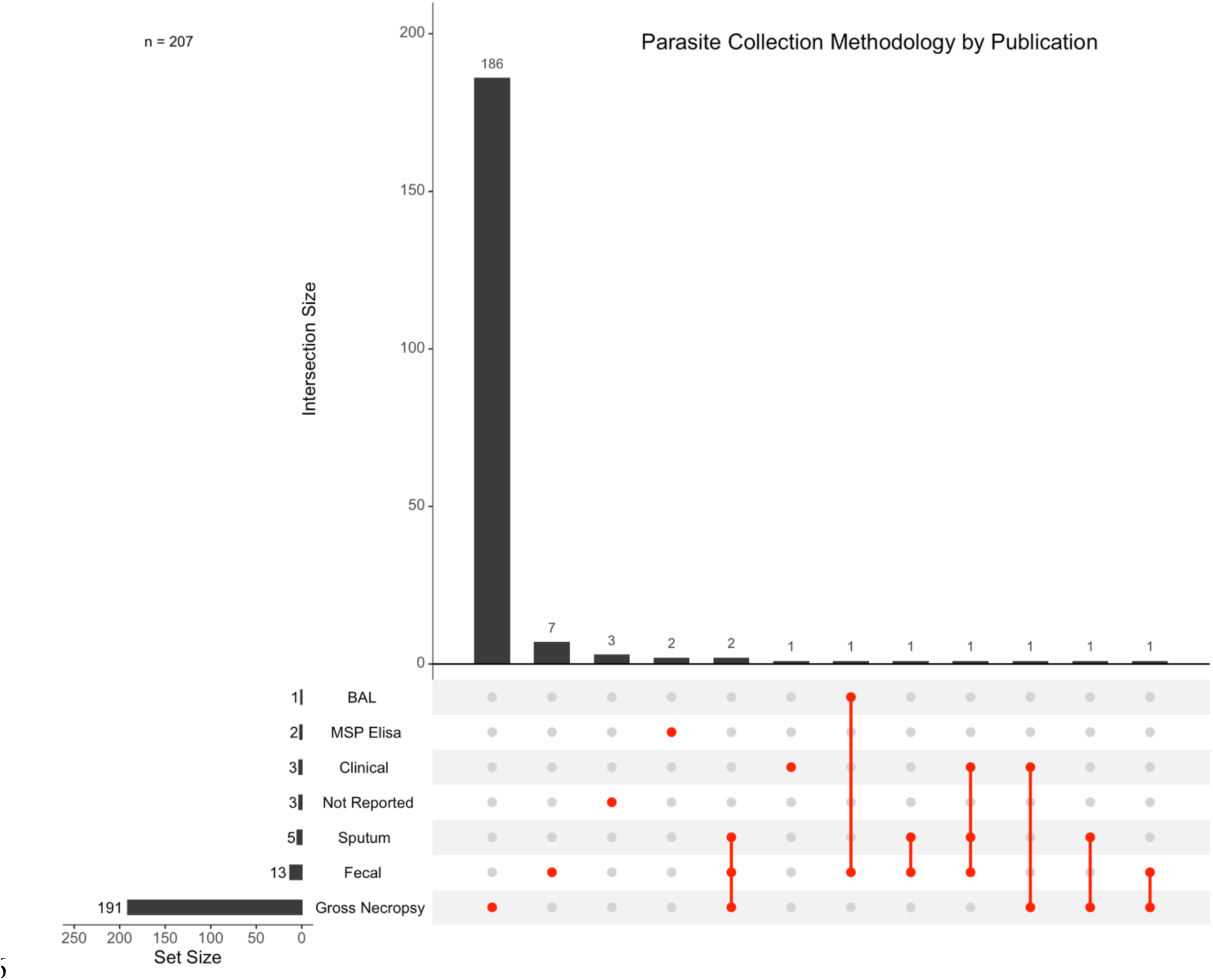
Parasite collection methodologies documented, with intersections between the methodologies shown to represent studies that utilize more than one methodology. Articles that did not report a collection methodology are designated as “Not Reported”.

A total of 175 host-metastrongyle relationships were reported in North America (including the commonly referenced Central America region), and 169 relationships were reported in Europe. North America and Europe are the most common locations reported for metastrongyle infections in marine mammals, followed by 54 relationships in South America, 24 in Asia, 23 in Oceania and 11 in Africa (Figure 3). Exactly half (50.00%, n=51/102) of all unique relationships are reported in North America, with 29.41% (n=30/102) of these relationships being reported exclusively in North America (Figure 4).

**Figure 3:**
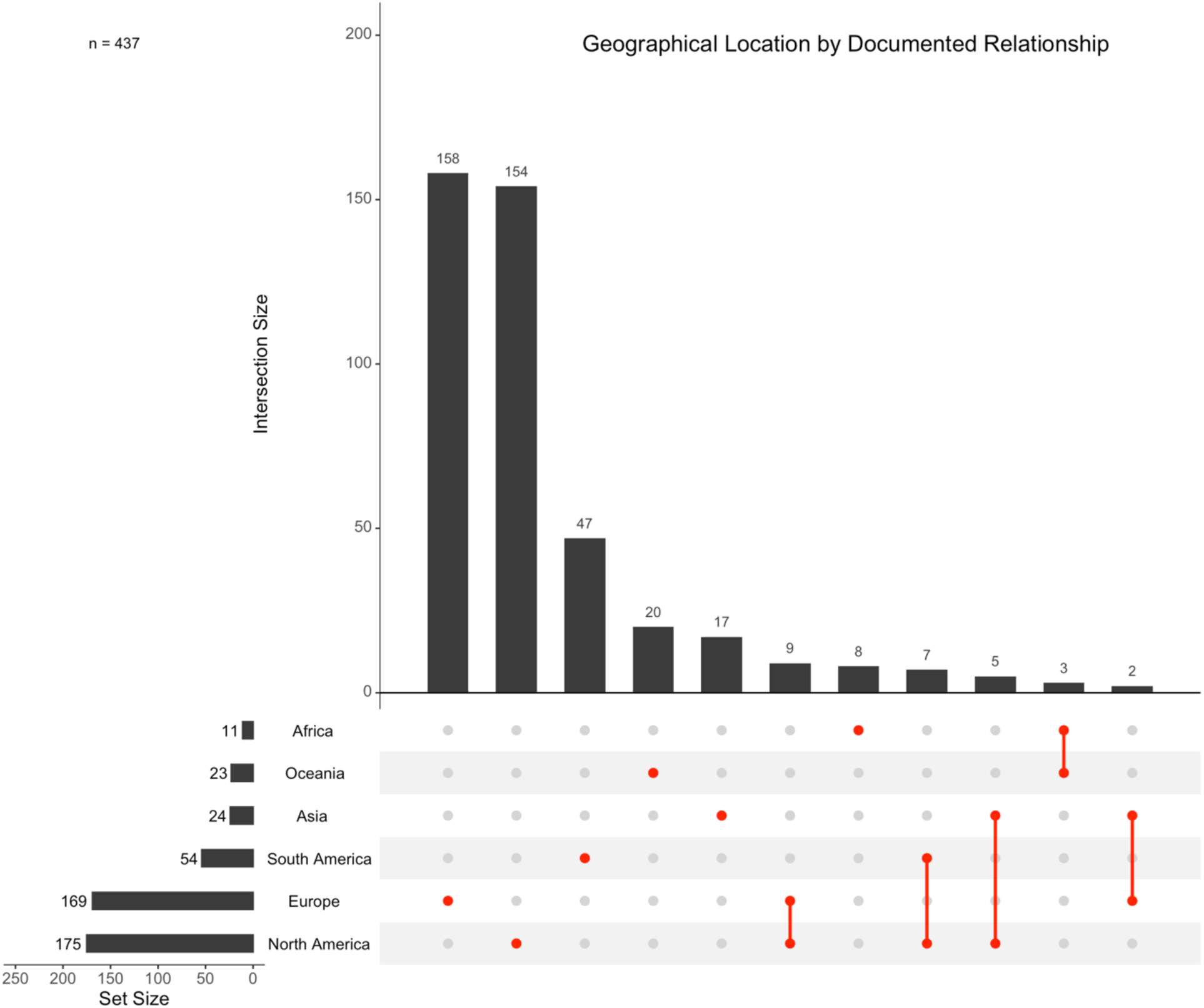
Geographical location(s), denoted by continent, of documented relationships between metastrongyles and marine mammals. Documented relationships include duplicates of unique relationships published independently. Intersections are shown for studies which report individual host species to parasite species relationships occurring in multiple locations.

**Figure 4:**
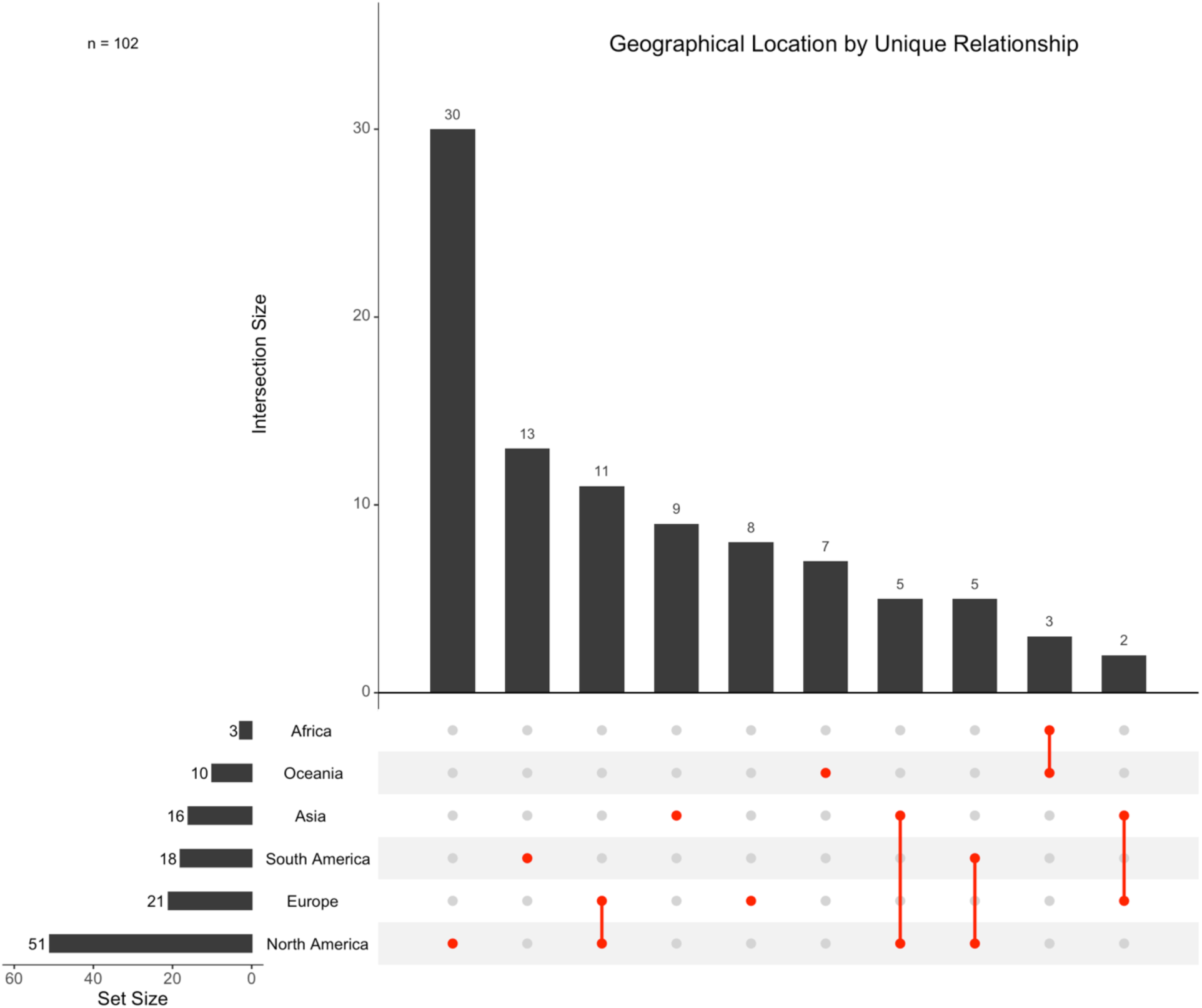
Geographical location(s), denoted by continent, of all unique relationships between metastrongyles and marine mammals. Intersections are shown for studies which report individual host species to parasite species relationships occurring in multiple locations.

### 3.2 Frequencies and diversities of metastrongyle infections in marine mammals

Pseudaliidae was the most reported metastrongyle family in marine mammals (n=331), followed by Filaroididae (n=90), and Crenosomatidae (n=58). At the host family level, Delphinidae (n=159), Phocoenidae (n=150), and Phocidae (n=107) are most frequently parasitized by metastrongyles. Further, metastrongyles from the Pseudaliidae family are reported in hosts from the Delphinidae (n=158), Phocoenidae (n=148), Monodontidae (n=15), Ziphiidae (n=3), Iniidae (n=2), Balaenopteridae (n=1), Kogiidae (n=1), and Physeteridae (n=1) families. Metastrongyles from the Crenosomatidae family are reported to parasitize hosts from the Phocidae (n=52), Otariidae (n=5), and Phocoenidae (n=1) families. Metastrongyles from the Filaroididae family are reported to parasitize hosts from the Phocidae (n=54), Otariidae (n=35), and Delphinidae (n=1) families.

*Otostrongylus circumlitus* (n=52) is the most reported metastrongyle species followed by *Stenurus minor* (n=42) and *Pseudalius inflexus* (n=31) (Supplementary Figure 1). The harbour porpoise (*Phocoena phocoena*) (n=118) is the most reported host, followed by harbour seal (*Phoca vitulina*) (n=41) and bottlenose dolphin (*Tursiops truncat*us) (n=23) (Supplementary Figure 2).

Before penalizing for literature and intra-study biases, the host species harbouring the largest number of metastrongyle species (parasite diversity) were: indo-pacific finless porpoise (*Neophocaena phocaenoides*) (n=8), short-beaked common dolphin (*Delphinus delphis*) (n=7), harbour porpoise (*Phocoena phocoena*) (n=7), and bottlenose dolphin (*Tursiops truncatus*) (n=7) (Supplementary Figure 3). After correction for publication bias, indo-pacific finless porpoise (*Neophocaena phocaenoides*) remained the most permissive host (Figure 5.). Before penalization, the metastrongyle species with the broader host diversity were: *Otostrongylus circumlitus* (n=13), *Stenurus mino*r (n=10), *Parafilaroides gymnurus* (n=9), *Halocercus brasiliensis* (n=8), and *Halocercus delphini* (n=8) (Supplementary Figure 4). After correction for publication bias, *Halocercus hyperdooni* emerged as the most generalist lungworm species, followed closely by *Otostrongylus circumlitus* and *Parafilaroides gymnurus* (Figure 6.).

**Figure 5:**
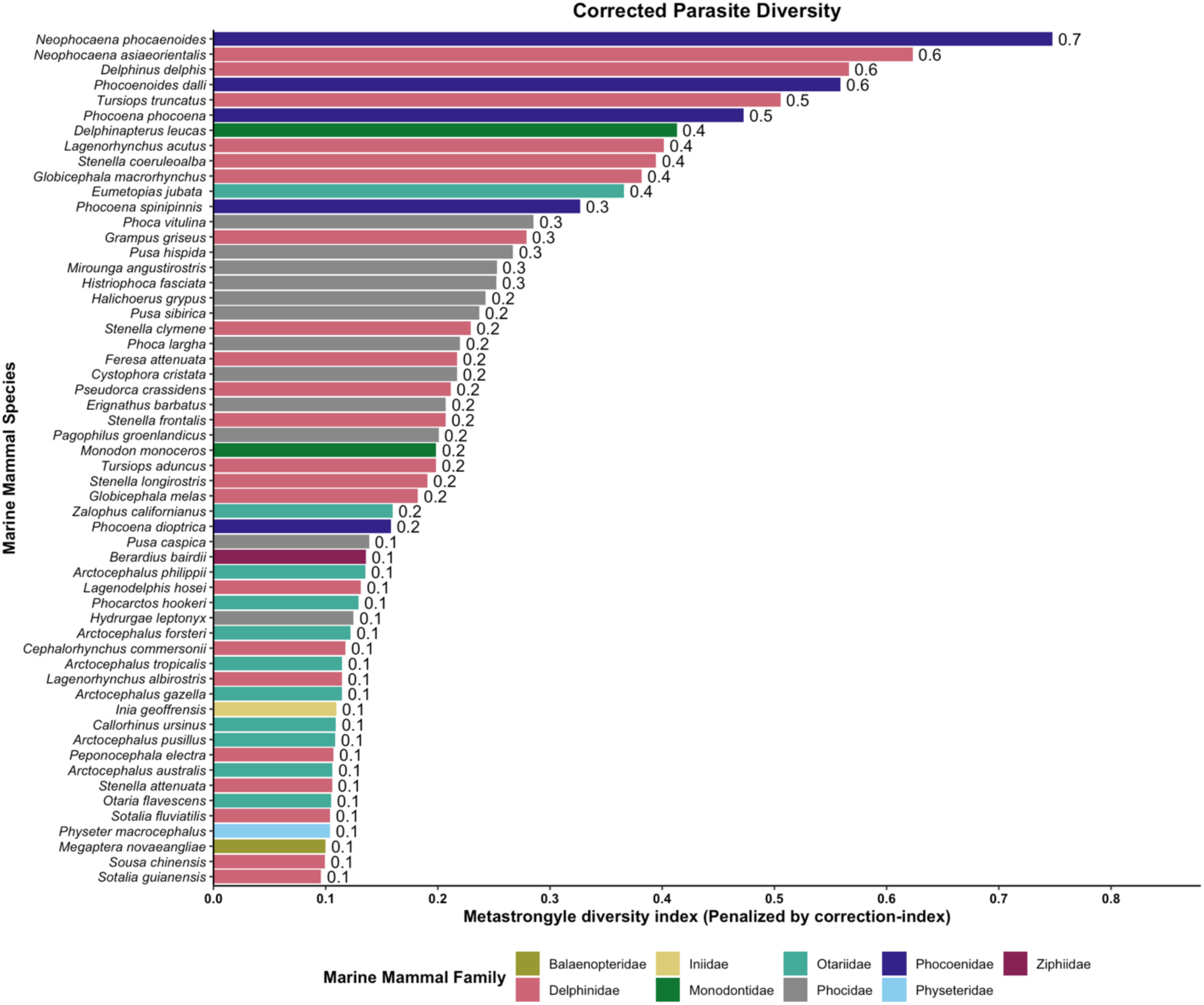
Parasite diversity for metastrongyles in marine mammals, corrected for publication bias (correction-index).

**Figure 6:**
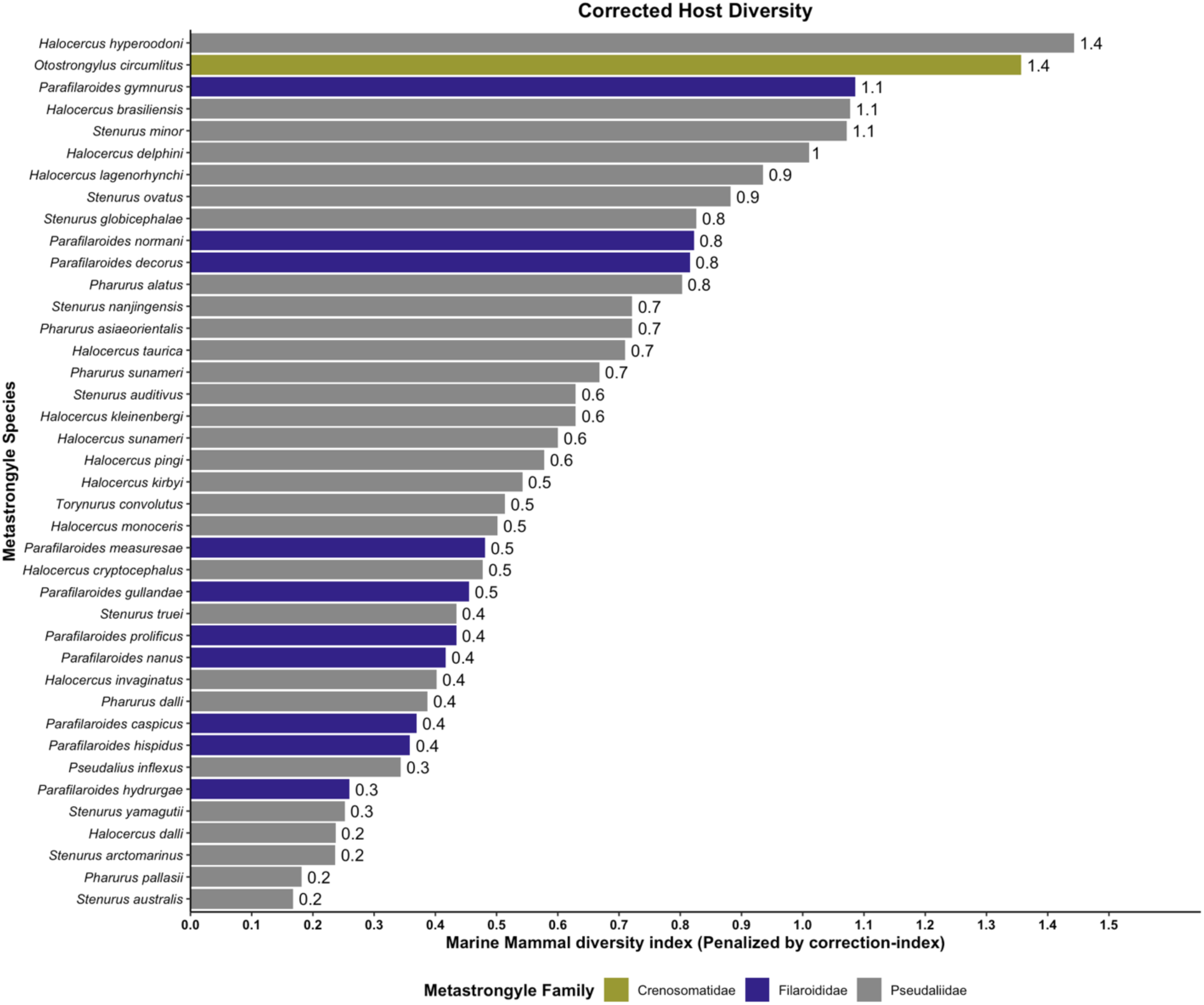
Host diversity for metastrongyles in marine mammals, corrected for publication bias (correction-index).

### 3.3 The impact of metastrongyle infections on marine mammals

Of the 481 host parasite relationships documented, 371 (77.13%) reported an anatomical location. Lungworms were found in the lower respiratory system in 82.48% (n=306/371) of cases, while parasites were found in the otic apparatus in 17.52% (n=65/371) of cases. In 11.59% (n=43/371) of cases, a single species of parasite parasitized an individual host in multiple anatomical locations (Figure 7).

**Figure 7:**
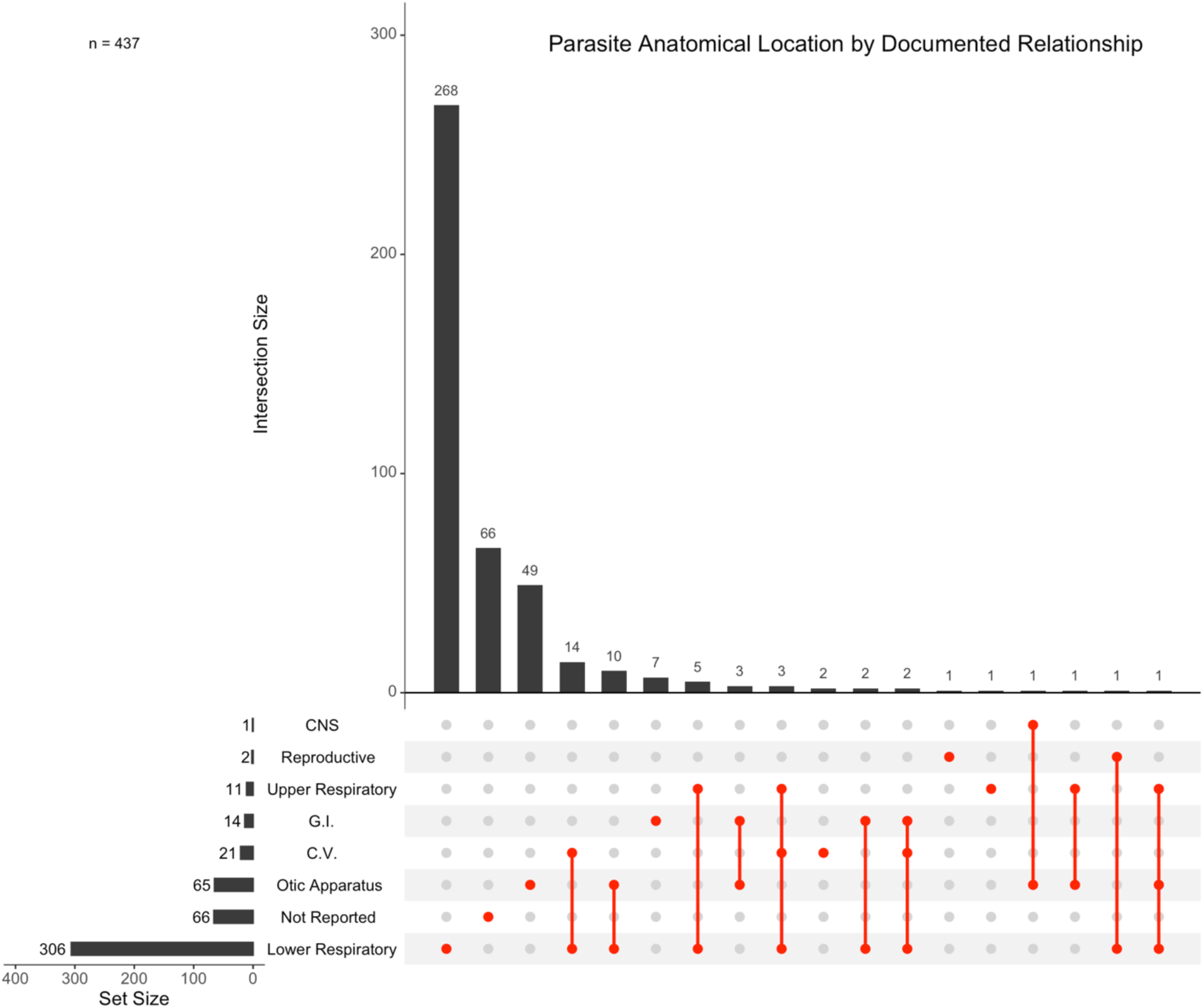
Anatomical location(s) of documented relationships between metastrongyles and marine mammals. Intersections are shown for studies which report individual host species to parasite species relationships occurring in multiple locations. Relationships that did not report an anatomical location are designated as “Not Reported”. (CNS= Central nervous system, G.I= Gastrointestinal tract, C.V= Cardiovascular system).

Out of the 201 unique host-parasite relationships assessed, 44.77% (n=90/201) mentioned mortality as “possible” or “potential” contributory cause of death and only 10.44% (n=21/201) studies considered metastrongyle infection as the primary cause of death. Of the 201 host-parasite relationships for which pathogenic effect was assessed, bronchopneumonia was described in 71.64% (n=144/201) of the relationships, indicating a high prevalence of bronchopneumonia in hosts infected with metastrongyles (Figure 8). Other reported pathogenic effects included edema (10.95%, n=22/201), vasculitis (7.46%, n=15/201), thrombosis (5.97%, n=12/201), and hemorrhage (5.97%, n=12/201). Lungworm parasitism resulted in no discernible pathogenic effect on the host in 16.42% (n=33/201) of the relationships.

**Figure 8:**
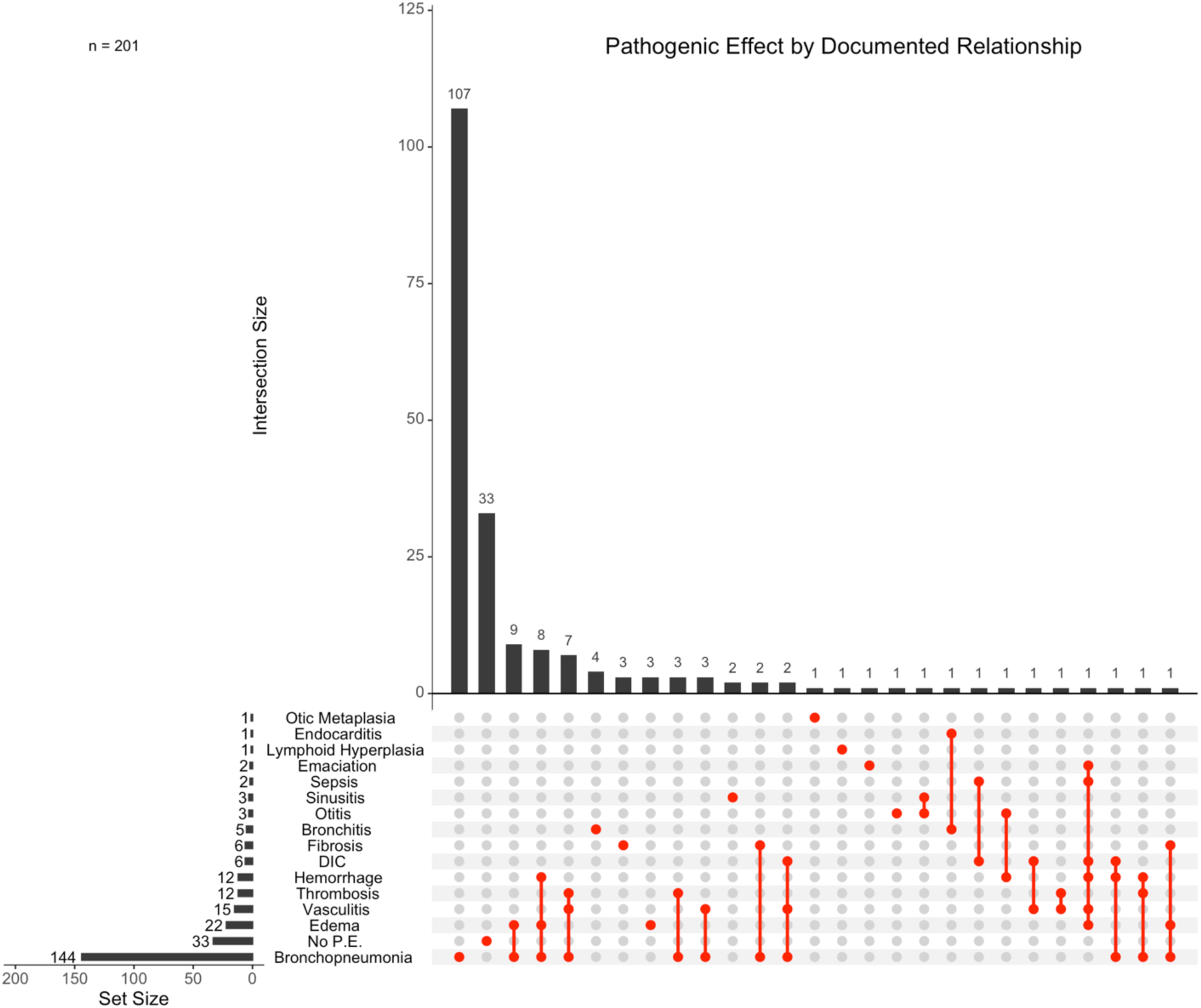
Pathogenic effect(s) of documented relationships between metastrongyles and marine mammals. Intersections are shown for studies which report individual host species to parasite species relationships causing multiple pathogenic effects. Relationships were designated as “No. P.E.” when pathogenic effect was assessed and was determined to be negligible. (DIC= Disseminated Intravascular Coagulation).

Pathogenic effect was assessed for 166 host-parasite relationships in the respiratory system, with respiratory specific pathology reported in 80.12% (n=133/166) of these cases. In contrast, 26 host-parasite relationships assessed pathogenic effect in the otic apparatus, with otic specific lesions reported in only 23.08% (n=6/26) of these relationships.

In the binomial GLMM that assessed pathogenic effect, host body size, corrected host diversity and corrected parasite diversity were all significant predictors of the presence of metastrongyle pathogenic effect in marine mammals, with a significant effect of the interaction between host and parasite diversity indexes (table 2). For respiratory specific effects, corrected parasite diversity and corrected host diversity main effects and their interaction were significant predictors of metastrongyle respiratory specific effect in marine mammals, but host body size was not a significant predictor of respiratory specific effect (table 3).

**Table 2:** Binomial generalized linear mixed effects model (GLMM) for presence of pathogenic effect. Presented are predictors estimates, standard error (SE), statistic (Z) and P values. The model used study ID as a random effect (total observations n=165, study ID n=89).

| Predictor | Estimate | SE | Z | P |
| --- | --- | --- | --- | --- |
| (Intercept) | -189.93 | 65.03 | -2.921 | 0.00349 |
| Corrected Parasite Diversity | 523.62 | 165.18 | 3.170 | 0.00152 |
| Corrected Host Diversity | 232.05 | 88.44 | 2.624 | 0.00869 |
| Host Body Size | 166.90 | 83.04 | 2.010 | 0.04444 |
| Corrected Parasite Diversity : Corrected Host Diversity | -625.90 | 250.44 | -2.499 | 0.01245 |
| Corrected Parasite Diversity : Host Body Size | -435.38 | 220.87 | -1.971 | 0.04870 |
| Corrected Host Diversity : Host Body Size | -189.10 | 131.22 | -1.441 | 0.14953 |
| Corrected Parasite Diversity : Corrected Host Diversity : Host Body Size | 581.44 | 438.86 | 1.325 | 0.18521 |

**Table 3:** Binomial generalized linear mixed effects model (GLMM) for respiratory pathogenic effects. Presented are predictors estimates, standard error (SE), statistic (Z) and P values. The model used study ID as a random effect (total observations n=134, study ID n=76).

| Predictor | Estimate | SE | Z | P |
| --- | --- | --- | --- | --- |
| (Intercept) | 0.09403 | 12.65860 | 0.007 | 0.99407 |
| Corrected Parasite Diversity | 88.93332 | 41.43432 | 2.146 | 0.03184 |
| Corrected Host Diversity | 74.10991 | 26.73963 | 2.772 | 0.00558 |
| Host Body Size | -18.39291 | 13.61594 | -1.351 | 0.17675 |
| Corrected Parasite Diversity : Corrected Host Diversity | -244.76967 | 87.16180 | -2.808 | 0.00498 |
| Corrected Parasite Diversity : Host Body Size | 8.03858 | 38.70274 | 0.208 | 0.83546 |
| Corrected Host Diversity : Host Body Size | -29.25975 | 24.54618 | -1.192 | 0.23325 |
| Corrected Parasite Diversity : Corrected Host Diversity : Host Body Size | 154.98741 | 93.04556 | 1.666 | 0.09577 |

## 4. Discussion and Conclusions

Metastrongyles are common helminth parasites of the respiratory and cardiovascular system of mammals. Given the importance of these body systems for the survival of diving animals, it was expected that the health impacts of metastrongyles would be particularly significant in marine mammals. Our findings confirm that pathogenic effects, mainly bronchopneumonia, are common in marine mammals infected with metastrongyles, although association of metastrongyle parasites and mortality is usually not definitive. Importantly, our review suggests that pathogenic host-parasite relationships are associated to hosts that can harbor a wider range of metastrongyle species (parasite diversity). From the parasite perspective, metastrongyle species with a broader host diversity tend to be more pathogenic. These results emphasize the importance of host and parasite diversity with regards to the virulence of metastrongyles in marine ecosystems.

Most parasitology studies in marine mammals have been conducted through the opportunistic collection and examination of stranded carcasses. This probably explains in part why in most cases metastrongyle infections are suggested or considered as a contributory cause of death but rarely confirmed as the primary cause of mortality. Determining cause of death in stranded marine mammals is challenging, amid the rapid rate of postmortem changes in these species and little background information for most cases (Arbelo *et al*., 2013; Alvarado-Rybak *et al*., 2020a). Additionally, access to carcasses is sometimes difficult (*e.g*., tides), limiting proper dissection techniques and tissue sampling. As with other host-parasite systems, the use of longitudinal assessments and experimental manipulation of infection could give new perspectives on the impact of metastrongyles in marine mammal health. Recently, the sampling of living species through the collection of feces and body fluids has been implemented in cetaceans (Williams *et al*., 2020). Further, the development of molecular techniques such as major sperm protein (MSP) ELISAs to test for serum antibodies to metastrongyles aid in illuminating metastrongyle seroprevalence (Ulrich *et al*., 2015, 2016). The continued use of approaches other than opportunistic necropsies may substantially contribute to the understanding of the drivers and consequences of metastrongyle infection in marine mammals.

As with many wildlife parasites (Haddaway and Watson, 2016; Seguel and Gottdenker, 2017), most studies on metastrongyles in marine mammals are conducted in North America and Europe, leading to overrepresentation of North American and European marine mammal and metastrongyle species in the literature. Therefore, cosmopolitan coastal marine mammals in these regions such as the harbor porpoise and the harbor seal along with their parasites were among the most assessed species. Despite our intended penalization for these study biases, our results should be interpreted with caution since more studies from different regions of the world and in cryptic and/or pelagic marine mammal species could influence inferences on the role of host and parasite factors driving pathogenic effects. For instance, PCBs and other persistent organic pollutants (POPs) are found in higher concentrations in marine mammals in more industrialized regions of North America and Europe (Fair *et al*., 2010; Jepson *et al*., 2016; Durante *et al*., 2016; Seguel *et al*., 2020a). These immunosuppressive pollutants are thought to increase susceptibility to metastrongyles, facilitating their pathogenic effects (Jepson *et al*., 2016). Therefore, more studies from regions of the world with lower POP concentrations or where new or underrepresented host-parasite relationships occur could modify our understanding of the health impacts of metastrongyles in marine mammals.

As in domestic and land mammals, bronchopneumonia is the most common pathogenic outcome of lungworm infection in marine mammals. The presence of lungworms in airways or pulmonary parenchyma causes tissue damage, eliciting an inflammatory response that also facilitates colonization by bacteria (Measures, 2001). Therefore, in most of the assessed studies, verminous bronchopneumonia was usually complicated by secondary bacterial infection. The impact of metastrongyles in other tissues such as blood vessels, heart or otic apparatus is probably more direct, since in these tissues bacteria colonization secondary to helminth infection is rare. In the cardiovascular system, the impact of lungworm infection can be serious with vasculitis, localized or systemic thrombosis and coagulopathy (Gulland *et al*., 1997b; Seguel *et al*., 2018b; Barnett *et al*., 2019b). In these cases, there is clear link between metastrongyle infection and mortality. Interestingly, invasion of the cardiovascular system is usually associated to metastrongyle species that are not the most common definitive host for that parasite species, highlighting the role of host specificity in the most pathogenic interactions between metastrongyles and marine mammals.

Parasite and host diversity increase the chances for a for a pathogenic outcome in metastrongyle-marine mammal relationships. This finding is supported by general hypotheses on parasite virulence that suggests that a parasite species with a greater diversity of potential hosts can evolve to be more pathogenic, as the sustainability of the parasite population is less dependent on the survival of individual hosts (Poulin, 2011; Schmid-Hempel, 2021). Similarly, a host species that harbors a greater diversity of parasites may see a greater propensity for pathogenic outcomes from individual parasite species due to an increase in the stress on the immune and respiratory systems resulting from co-infections (Seguel *et al*., 2023). Our results also suggest that larger hosts are more likely to sustain pathogenic outcomes because of metastrongyle parasitism. Since marine metastrongyles are transmitted through intermediate hosts that represent food items for marine mammals (Dailey, 1970a), this result could indicate that increased food consumption leads to a greater larvae intake and subsequently a higher parasite burden. Similarly, a larger host body size could correlate with access to different prey items, contributing to a greater diversity of metastrongyle infections.

A comprehensive understanding of the drivers of the negative physiological effects associated with metastrongyle parasitism in marine mammal hosts is challenged by a presumably strong bias towards reporting only positive findings. The available literature clearly elucidates a multitude of host-parasite relationships in which metastrongyles produce a definable pathological effect, howbeit it is inadequate in clarifying why these effects are observed in some species or populations and not in others. However, general host and parasite traits associated with pathogenic outcomes are like those of other helminth and microbial parasites, including host permissiveness of infection by multiple species and parasite capacity to infect multiple host species. In this context, the rapidly changing marine environment with generalized loss of biomass could favor parasites and hosts with a diversified trophic network, potentially selecting for more pathogenic relationships between metastrongyles and their marine mammal hosts.

## 5. Supplementary Data

Supplementary figures are submitted as electronic supplementary files. Review protocol not registered.

## 6. Financial Support

This work was supported by the University of Guelph President’s Research Assistantship, which was awarded to J.F. as a component of his University of Guelph President’s Scholarship funding. The Author’s declare no competing interest.

## Notes

### Competing Interest Statement

The authors have declared no competing interest.

### Summary of Updates

Formatting revisions.

## References

Abollo, E., López, A., Gestal, C., Benavente, P. and Pascual, S. (1998). Macroparasites in cetaceans stranded on the northwestern Spanish Atlantic coast. Diseases of Aquatic Organisms 32, 227–231. doi: 10.3354/dao032227.

Aguilar-Aguilar, R., Delgado-Estrella, A. and Moreno-Navarrete, R. (2010). New host report for nematodes in a stranded short-snouted spinner dolphin Stenella clymene (Cetacea: Delphinidae) from the Mexican Caribbean coast. Helminthologia 47, 136–138. doi: 10.2478/s11687-010-0020-0.

Alvarado-Rybak, M., Toro, F., Abarca, P., Paredes, E., Español-Jiménez, S. and Seguel, M. (2020a). Pathological Findings in Cetaceans Sporadically Stranded Along the Chilean Coast. Frontiers in Marine Science 7, 1–10. doi: 10.3389/fmars.2020.00684.

Alvarado-Rybak, M., Toro, F., Abarca, P., Paredes, E., Español-Jiménez, S. and Seguel, M. (2020b). Pathological Findings in Cetaceans Sporadically Stranded Along the Chilean Coast. Frontiers in Marine Science 7, 684. doi: 10.3389/fmars.2020.00684.

Andersen, S. H. (1974). A Typical Case History of the Net-Caught Harbour Porpose. Aquatic Mammals 2, 1–6.

Antonelis, G. A., Melin, S. R. and Bukhtiyarov, Y. A. (1994). Early Spring Feeding Habits of Bearded Seals (Erignathus Barbatus) in the Central Bering Sea, 1981. ARCTIC 47, 74–79. doi: 10.14430/arctic1274.

Arbelo, M., De Los Monteros, A. E., Herráez, P., Andrada, M., Sierra, E., Rodríguez, F., Jepson, P. D. and Fernández, A. (2013). Pathology and causes of death of stranded cetaceans in the canary Islands (1999-2005). Diseases of Aquatic Organisms 103, 87–99. doi: 10.3354/dao02558.

Arnold, P. W. and Gaskin, D. E. (1975). Lungworms (Metastrongyloidea: Pseudaliidae) of harbor porpoise Phocoena phocoena (L. 1758). Canadian Journal of Zoology 53, 713– 735. doi: 10.1139/z75-087.

Aytemiz, I., Dede, A., Danyer, E. and Tonay, A. M. (2012). Morphological identification of parasites found in the stomach contents of bycaught striped dolphins (Stenella coeruleoalba) from Turkish Eastern Mediterranean Sea coast. Journal of the Black Sea / Mediterranean Environment 18, 238–245.

Babero, B. B. and Thomas, L. J. (1960). A Record of Pharurus oserkaiae (Skrjabin, 1942) in an Alaskan Whale. The Journal of Parasitology 46, 726. doi: 10.2307/3275519.

Bakbergenuly, I. and Kulinskaya, E. (2018). Meta-analysis of binary outcomes via generalized linear mixed models: a simulation study. BMC Medical Research Methodology 18, 70. doi: 10.1186/s12874-018-0531-9.

Baker, J. R. (1987). Causes of mortality and morbidity in wild juvenile and adult grey seals (Halichoerus grypus). British Veterinary Journal 143, 203–220. doi: 10.1016/0007-1935(87)90083-2.

Balbuena, J. A., Aspholm, P. E., Andersen, K. I. and Bjørge, A. (1994). Lung-worms (Nematoda: Pseudaliidae) of harbour porpoises (*Phocoena phocoena*) in Norwegian waters: patterns of colonization. Parasitology 108, 343–349. doi: 10.1017/S0031182000076186.

Barnett, J. E. F., Bexton, S., Fraija-Fernández, N., Chooneea, D. and Wessels, M. E. (2019a). Novel Pulmonary Vasculitis with Splendore–Hoeppli Reaction in Grey Seals (Halichoerus grypus) Associated with Otostrongylus circumlitus Infection. Journal of Comparative Pathology 173, 83–91. doi: 10.1016/j.jcpa.2019.10.009.

Barnett, J. E. F., Bexton, S., Fraija-Fernández, N., Chooneea, D. and Wessels, M. E. (2019b). Novel Pulmonary Vasculitis with Splendore–Hoeppli Reaction in Grey Seals (Halichoerus grypus) Associated with Otostrongylus circumlitus Infection. Journal of Comparative Pathology 173, 83–91. doi: 10.1016/j.jcpa.2019.10.009.

Barnett, J., Allen, R., Astley, K., Whitehouse, F. and Wessels, M. (2021). Pathology of grey seals Halichoerus grypus in southwest England including pups in early rehabilitation. Diseases of Aquatic Organisms 145, 35–50. doi: 10.3354/dao03600.

Baylis, H. A. (1932). A list of worms parasitic in Cetacea. Discovery Reports 6, 393–418.

Baylis, H. A. and Daubney, R. (1925). A Revision of the Lung-Worms of Cetacea. Parasitology 17, 201–215. doi: 10.1017/S0031182000004595.

Bergeron, E., Huot, J. and Measures, L. N. (1997a). Experimental transmission of Otostrongylus circumlitus (Railliet, 1899) (Metastrongyloidea: Crenosomatidae), a lungworm of seals in eastern arctic Canada. Canadian Journal of Zoology 75, 1364–1371. doi: 10.1139/z97-762.

Bergeron, E., Measures, L. N. and Huot, J. (1997b). Lungworm (Otostrongylus circumlitus) infections in ringed seals (Phoca hispida) from eastern Arctic Canada. Canadian Journal of Fisheries and Aquatic Sciences 54, 2443–2448. doi: 10.1139/f97-153.

Beverley-Burton, M. (1978). Helminths of the Alimentary Tract from a Stranded Herd of the Atlantic White-Sided Dolphin, *Lagenorhynchus acutus*. Journal of the Fisheries Research Board of Canada 35, 1356–1359. doi: 10.1139/f78-211.

Birincioğlu, S. S., Aypak, S., Avci, H., Birincioğlu, B., İPek, E. and Akkoç, A. N. (2017). Şişe Burunlu Bir Yunusta (Tursiops truncatus) Patolojik ve Parazitolojik İncelemeler. Kafkas Universitesi Veteriner Fakultesi Dergisi. doi: 10.9775/kvfd.2017.18016.

Birkun, A. A. Jr. and Krivokhizhin, S. V. (1996). Present state and causes of the Black Sea cetacean populations suppression. Parts I and II. Vestnik Zoologii 3, 36–42.

Borgsteede, F. H. M., Bus, H. G. J., Verplanke, J. A. W. and van Burg, W. P. J. (1991). Endoparasitic helminths of the harbour seal, Phoca vitulina, in the Netherlands. Netherlands Journal of Sea Research 28, 247–250. doi: 10.1016/0077-7579(91)90022-S.

Bowie, J. Y. (1984). Parasites from an Atlantic bottle-nose dolphin (*Tursiops truncatus*), and a revised checklist of parasites of this host. New Zealand Journal of Zoology 11, 395–398. doi: 10.1080/03014223.1984.10428253.

Brooks, M. E., Kristensen, K., van Benthem, K. J., Magnusson, A., Berg, C. W., Nielsen, A., Skaug, H. J., Mächler, M. and Bolker, B. M. (2017). glmmTMB balances speed and flexibility among packages for zero-inflated generalized linear mixed modeling. R Journal 9, 378–400. doi: 10.32614/rj-2017-066.

Brosens, L., Jauniaux, T., Siebert, U., Benke, H. and Coignoul, F. (1996). Observations on the helminths of harbour porpoises (Phocoena phocoena) and common guillemots (Uria aalge) from the Belgian and German coasts. Veterinary Record 139, 254–257. doi: 10.1136/vr.139.11.254.

Burek-Huntington, K., Dushane, J., Goertz, C., Measures, L., Romero, C. and Raverty, S. (2015). Morbidity and mortality in stranded Cook Inlet beluga whales Delphinapterus leucas. Diseases of Aquatic Organisms 114, 45–60. doi: 10.3354/dao02839.

Cannon, L. R. G. (1977). Some aspects of the biology of Peponocephala electra (Cetacea: Delphinidae). II. Parasites. Marine and Freshwater Research 28, 717–722.

Carvalho, V. L., Bevilaqua, C. M. L., Iñiguez, A. M., Mathews-Cascon, H., Ribeiro, F. B., Pessoa, L. M. B., de Meirelles, A. C. O., Borges, J. C. G., Marigo, J., Soares, L. and de Lima Silva, F. J. (2010). Metazoan parasites of cetaceans off the northeastern coast of Brazil. Veterinary Parasitology 173, 116–122. doi: 10.1016/j.vetpar.2010.06.023.

Carvalho Demarque, I. de O., Rodrigues de Oliveira, F. C., da Silveira, L. S., Barbosa, L. A. and Ederli, N. B. (2020). The Lungworm, Halocercus brasiliensis (Nematoda: Pseudaliidae), from Guiana Dolphins Sotalia guianensis from Brazil with Pathological Findings. Journal of Parasitology 106, 254. doi: 10.1645/19-77.

Claussen, D., Strauss, V., Ising, S., Jäger, M., Schnieder, T. and Stoye, M. (1991). The Helminth Fauna from the Common Seal (Phoca vitulina vitulina, Linné, 1758) of the Wadden Sea in Lower Saxony*: Part 2: Nematodes. Journal of Veterinary Medicine, Series B 38, 649–656. doi: 10.1111/j.1439-0450.1991.tb00924.x.

Colegrove, K. M., Greig, D. J. and Gulland, F. M. D. (2005). Causes of Live Strandings of Northern Elephant Seals (Mirounga angustirostris) and Pacific Harbor Seals (Phoca vitulina) Along the Central California Coast, 1992-2001. Aquatic Mammals 31, 1–10. doi: 10.1578/AM.31.1.2005.1.

Colón-Llavina, M. M., Mattiucci, S., Nascetti, G., Harvey, J. T., Williams, E. H. and Mignucci-Giannoni, A. A. (2019). Some Metazoan Parasites from Marine Mammals Stranded in California. Pacific Science 73, 461. doi: 10.2984/73.4.3.

Conlogue, G. J., Ogden, J. A. and Foreyt, W. J. (1985). Parasites of the Dall’s porpoise (Phocoenoides dalli True). Journal of Wildlife Diseases 21, 160–166. doi: 10.7589/0090-3558-21.2.160.

Corcuera, J., Monzon, F., Anguilar, A., Borrell, A. and Raga, J. A. (1995). Life history data, organochlorine pollutants and parasites from eight Burmeister’s porpoises, Phocoena spinipinnis, caught in northern Argentine waters. Report of the International Whaling Commission.

Cowan, D. F. (1967). Helminth Parasites of the Pilot Whale Globicephala melaena (Traill 1809). The Journal of Parasitology 53, 166. doi: 10.2307/3276641.

Dagleish, M. P., Barley, J., Finlayson, J., Reid, R. J. and Foster, G. (2008). Brucella ceti Associated Pathology in the Testicle of a Harbour Porpoise (Phocoena phocoena). Journal of Comparative Pathology 139, 54–59. doi: 10.1016/j.jcpa.2008.03.004.

Dailey, M. D. (1970a). The Transmission of Parafilaroides decorus (Nematoda : Metastrongyloidea) in the California Sea Lion (Zalophus calif ornianus) 1. Proc. Helm. Soc. Wash. 37, 215–222.

Dailey, M. D. (1970b). The Transmission of Parafilaroides decorus (Nematoda: Metastrongyloidea) in the California Sea Lion. Proceedings of the Helminthological Society of Washington 37, 215–222.

Dailey, M. D. (2006a). Restoration of Parafilaroides (Dougherty, 1946) (Nematoda: Metastrongyloidea) with description of two new species from pinnipeds of eastern central Pacific. The Journal of Parasitology 92, 589–594. doi: 10.1645/GE-3525.1.

Dailey, M. D. (2006b). Restoration of Parafilaroides (Dougherty, 1946) (Nematoda: Metastrongyloidea) with Description of Two New Species from Pinnipeds of Eastern Central Pacific. Journal of Parasitology 92, 589–594. doi: 10.1645/GE-3525.1.

Dailey, M. D. (2009). A New Species of Parafilaroides (Nematoda: Filaroididae) in Three Species of Fur Seals (Carnivora: Otariidae) From the Southern Hemisphere. Journal of Parasitology 95, 156–159. doi: 10.1645/GE-1521.1.

Dailey, M. D. and Brownell, R. L. Jr. (1972). A checklist of marine mammal parasites. In Mammals of the sea: Biology and medicine, SH Ridgway (ed.). Charles C Thomas, Springfield, Illinois, pp. 528–589.

Dailey, M. D. and Perrin, W. F. (1973). Helminth parasities of porpoises of the genus Stenella in the eastern tropical pacific, with descriptions of two new species: Mastigonema stenellae gen. et. sp. n. (Nematoda: spiruroidea) and Zalophotrema pacificum sp. n. (Trematoda: digenea). Fishery Bulletin 71, 455–471.

Dailey, M. and Stroud, R. (1978). Parasites and associated pathology observed in cetaceans stranded along the Oregon coast. Journal of Wildlife Diseases 14, 503–511. doi: 10.7589/0090-3558-14.4.503.

Dailey, M. D. and Walker, W. A. (1978). Parasitism as a Factor (?) in Single Strandings of Southern California Cetaceans. The Journal of Parasitology 64, 593. doi: 10.2307/3279939.

Dailey, M., Walsh, M., Odell, D. and Campbell, T. (1991a). Evidence of Prenatal Infection in the Bottlenose Dolphin (Tursiops truncates) with the Lungworm Halocercus lagenorhynchi (Nematoda: Pseudaliidae). Journal of Wildlife Diseases 27, 164–165. doi: 10.7589/0090-3558-27.1.164.

Dailey, M., Walsh, M., Odell, D. and Campbell, T. (1991b). Evidence of prenatal infection in the bottlenose dolphin (Tursiops truncatus) with the lungworm Halocercus lagenorhynchi (Nematoda: Pseudaliidae). Journal of Wildlife Diseases 27, 164–165. doi: 10.7589/0090-3558-27.1.164.

Davison, N. J., Simpson, V. R., Chappell, S., Monies, R. J., Stubberfield, E. J., Koylass, M., Quinney, S., Deaville, R., Whatmore, A. M. and Jepson, P. D. (2010a). Prevalence of a host-adapted group B Salmonella enterica in harbour porpoises (Phocoena phocoena) from the south-west coast of England. Veterinary Record 167, 173–176. doi: 10.1136/vr.c3760.

Davison, N., Barnett, J., Rule, B., Chappell, S. and Wise, G. (2010b). Group B Salmonella in lungworms from a harbour porpoise (Phocoena phocoena). Veterinary Record 167, 351–352. doi: 10.1136/vr.c4495.

Davison, N. J., Barnett, J. E. F., Perrett, L. L., Dawson, C. E., Perkins, M. W., Deaville, R. C. and Jepson, P. D. (2013). Meningoencephalitis and Arthritis Associated with Brucella ceti in a Short-beaked Common Dolphin (Delphinus delphis). Journal of Wildlife Diseases 49, 632–636. doi: 10.7589/2012-06-165.

Dawson, C. E., Perrett, L. L., Stubberfield, E. J., Stack, J. A., Farrelly, S. S. J., Cooley, W. A., Davison, N. J. and Quinney, S. (2008). Isolation and characterization of Brucella from the lungworms of a Harbour porpoise (Phocoena phocoena). Journal of Wildlife Diseases 44, 237–246. doi: 10.7589/0090-3558-44.2.237.

de Castro, R. L., Vales, D. G., Degrati, M., García, N., Fernández, M. and Crespo, E. A. (2014). First record of cestode cysts of Phyllobothrium delphini (Phyllobothriidae) from dusky dolphins (Lagenorhynchus obscurus) off Argentine coast. Hidrobiológica 24, 307– 310.

Delyamure, S. L. (1968). Helminthofauna of Marine Mammals: Ecology and Phylogeny. ed. Skri͡abin, K. I. Israel Program for Scientific Translations.

Delyamure, S., Yurakhno, M., Popov, V. N., Shults, L. M. and Fay, F. H. (1984). Helminthological comparison of subpopulations of Bering Sea spotted seals, Phoca largha Pallas. Soviet-American cooperative research on marine mammals 1, 61–65.

Dennison, S., Gulland, F., Haulena, M., De Morais, H. and Colegrove, K. (2007). Urate nephrolithiasis in a northern elephant seal (Mirounga angustirostris) and a California sea lion (Zalophus californianus). Journal of Zoo and Wildlife Medicine 38, 114–120. doi: 10.1638/05-121.1.

Di Beneditto, A. P. M. and Ramos, R. M. A. (2004). Biology of the marine tucuxi dolphin (Sotalia fluviatilis) in south-eastern Brazil. Journal of the Marine Biological Association of the United Kingdom 84, 1245–1250. doi: 10.1017/S0025315404010744h.

DÍaz-Delgado, J., Arbelo, M., Sacchini, S., Quesada-Canales, Ó., Andrada, M., Rivero, M. and Fernández, A. (2012). Pulmonary Angiomatosis and Hemangioma in Common Dolphins (*Delphinus delphis*) Stranded in Canary Islands. Journal of Veterinary Medical Science 74, 1063–1066. doi: 10.1292/jvms.11-0573.

Domiciano, I. G., Domit, C., Broadhurst, M. K., Koch, M. S. and Bracarense, A. P. F. R. L. (2016). Assessing Disease and Mortality among Small Cetaceans Stranded at a World Heritage Site in Southern Brazil. PLOS ONE 11, e0149295. doi: 10.1371/journal.pone.0149295.

Dougherty, E. C. (1943). Notes on the lungworms of porpoises and their occurrence on the California coast. Proceedings of the Helminthological Society of Washington 10, 16–22.

Dougherty, E. C. (1944). The lungworms (Nematoda: Pseudalidae) of Odontoceti. Part I. Parasitology 36, 80–94.

Dougherty, E. C. and Herman, C. M. (1947). New species of the genus Parafilaroides Dougherty, 1946 (Nematoda: Metastrongylidae) from sea-lions, with a list of the lungworms of the Pinnipedia. Proceedings of the Helminthological Society of Washington 14, 77–87.

Dougnac, C. and Fredes, F. (2012). Identificación de fauna endoparasitaria en cetáceos de Tierra del Fuego: Nuevos registros de helmintos para la zona y para odontocetos de la región. Editorial Academica Espanola.

Duignan, P. J. (2000). Diseases of cetaceans and pinnipeds. In Marine wildlife ; the Fabian Fay course for veterinarians ; proceedings 335, 4-8 September 2000, pp. 419–447. Univ. of Sydney, Sydney.

Dunn, J. L. and Wolke, R. E. (1976). Dipetalonema spirocauda infection in the atlantic harbour seal (Phoca vitulina concolor). Journal of Wildlife Diseases 12, 531–538. doi: 10.7589/0090-3558-12.4.531.

Durante, C. A., Santos-Neto, E. B., Azevedo, A., Crespo, E. A. and Lailson-Brito, J. (2016). POPs in the South Latin America: Bioaccumulation of DDT, PCB, HCB, HCH and Mirex in blubber of common dolphin (Delphinus delphis) and Fraser’s dolphin (Lagenodelphis hosei) from Argentina. Science of The Total Environment 572, 352–360. doi: 10.1016/j.scitotenv.2016.07.176.

Ebmer, D., Navarrete, M. J., Muñoz, P., Flores, L. M., Gärtner, U., Brabec, J., Poppert, S., Taubert, A. and Hermosilla, C. (2020). Anthropozoonotic Parasites Circulating in Synanthropic and Pacific Colonies of South American Sea Lions (Otaria flavescens): Non-invasive Techniques Data and a Review of the Literature. Frontiers in Marine Science 7, 543829. doi: 10.3389/fmars.2020.543829.

Echenique, J., Pereira, E., Prado, J., Schild, A. L. and Valente, A. L. (2020). New host and geographical records for Parafilaroides normani (Nematoda: Filaroididae) Dailey, 2009 in South American fur seal, Arctocephalus australis, from southern Brazil. Journal of Helminthology 94, 1–4. doi: 10.1017/S0022149X20000826.

Elson-Riggins, J. G., Al-Banna, L., Platzer, E. G. and Kaloshian, I. (2001). Characterization of Otostrongylus circumlitis from pacific harbor and northern elephant seals. Journal of Parasitology 87, 73–78.

Elson-Riggins, J. G., Riggins, S. A., Gulland, F. M. D. and Platzer, E. G. (2004). Immunoglobulin responses of northern elephant and Pacific harbor seals naturally infected with Otostrongylus circumlitus. Journal of Wildlife Diseases 40, 466–475. doi: 10.7589/0090-3558-40.3.466.

Elson-Riggins, J. G., Gibbons, L. M., Van Liere, D. W., Zinkstok, E. W., Blake, D. P., Alegre, F., Spittle, H., Brakefield, P. M., Udo de Haes, H. A. and Osinga, N. (2020). Surprisingly long body length of the lungworm Parafilaroides gymnurus from common seals of the Dutch North Sea. Parasitology Research 119, 1803–1817. doi: 10.1007/s00436-020-06675-7.

Evans, P. (1991). Whales, dolphins and porpoises: order Cetacea. In The handbook of British mammals, pp. 299–350. Blackwell Oxford.

Evans, R. H. (2011). Segniliparus rugosus–associated Bronchiolitis in California Sea Lion. Emerging Infectious Diseases 17, 311–312. doi: 10.3201/eid1702.101511.

Ewing, R. Y., Rotstein, D. S., McLellan, W. A., Costidis, A. M., Lovewell, G., Schaefer, A. M., Romero, C. H. and Bossart, G. D. (2020). Macroscopic and Histopathologic Findings From a Mass Stranding of Rough-Toothed Dolphins (Steno bredanensis) in 2005 on Marathon Key, Florida, USA. Frontiers in Veterinary Science 7, 572. doi: 10.3389/fvets.2020.00572.

Ezenwa, V. O., Price, S. A., Altizer, S., Vitone, N. D. and Cook, K. C. (2006). Host traits and parasite species richness in even and odd-toed hoofed mammals, Artiodactyla and Perissodactyla. Oikos 115, 526–536. doi: 10.1111/j.2006.0030-1299.15186.x.

Fair, P. A., Adams, J., Mitchum, G., Hulsey, T. C., Reif, J. S., Houde, M., Muir, D., Wirth, E., Wetzel, D., Zolman, E., McFee, W. and Bossart, G. D. (2010). Contaminant blubber burdens in Atlantic bottlenose dolphins (Tursiops truncatus) from two southeastern US estuarine areas: Concentrations and patterns of PCBs, pesticides, PBDEs, PFCs, and PAHs. Science of the Total Environment 408, 1577–1597. doi: 10.1016/j.scitotenv.2009.12.021.

Faulkner, J. (1995). Study of the cranial sinus nematode Stenurus minor, Metastrongyloidea, in the harbour porpoise, Phocoena phocoena.

Faulkner, J., Measures, L. N. and Whoriskey, F. G. (1998). Stenurus minor (Metastrongyloidea: Pseudaliidae) infections of the cranial sinuses of the harbour porpoise, Phocoena phocoena. Canadian Journal of Zoology 76, 1209–1216.

Fauquier, D., Gulland, F., Haulena, M. and Spraker, T. (2003). Biliary Adenocarcinoma in a Stranded Northern Elephant Seal (Mirounga angustirostris). Journal of Wildlife Diseases 39, 723–726. doi: 10.7589/0090-3558-39.3.723.

Fauquier, D., Gulland, F., Haulena, M., Dailey, M., Rietcheck, R. L. and Lipscomb, T. P. (2004). Meningoencephalitis in Two Stranded California Sea Lions (Zalophus californianus) Caused by Aberrant Trematode Migration. Journal of Wildlife Diseases 40, 816–819. doi: 10.7589/0090-3558-40.4.816.

Fauquier, D., Kinsel, M., Dailey, M., Sutton, G., Stolen, M., Wells, R. and Gulland, F. (2009). Prevalence and pathology of lungworm infection in bottlenose dolphins Tursiops truncatus from southwest Florida. Diseases of Aquatic Organisms 88, 85–90. doi: 10.3354/dao02095.

Fauquier, D. A., Kinsel, M. J., Dailey, M. D., Sutton, G. E., Stolen, M. K., Wells, R. S. and Gulland, F. M. D. (2010). Prevalence and pathology of lungworm infection in bottlenose dolphins Tursiops truncatus from southwest Florida. Diseases of Aquatic Organisms 88, 85–90. doi: 10.3354/dao02095.

Fenton, H., Daoust, P., Forzán, M., Vanderstichel, R., Ford, J., Spaven, L., Lair, S. and Raverty, S. (2017). Causes of mortality of harbor porpoises Phocoena phocoena along the Atlantic and Pacific coasts of Canada. Diseases of Aquatic Organisms 122, 171–183. doi: 10.3354/dao03080.

Fernández, M., Agustí, C., Aznar, F. and Raga, J. (2003). Gastrointestinal helminths of Risso’s dolphin Grampus griseus from the Western Mediterranean. Diseases of Aquatic Organisms 55, 73–76. doi: 10.3354/dao055073.

Fleischman, R. W. and Squire, R. A. (1970). Verminous Pneumonia in the California Sea Lion (Zalophus californianus). Pathologia veterinaria 7, 89–101. doi: 10.1177/030098587000700201.

Forbes, A. B. (2021). Lungworm infections in cattle. In Parasites of cattle and sheep: a practical guide to their biology and control, pp. 116–138. CAB International, UK doi: 10.1079/9781789245158.0116.

Forrester, D. J., Odell, D. K., Thompson, N. P. and White, J. R. (1980). Morphometrics, Parasites, and Chlorinated Hydrocarbon Residues of Pygmy Killer Whales from Florida. Journal of Mammalogy 61, 356–360. doi: 10.2307/1380067.

Fouchier, R. A. M., Bestebroer, T. M., Martina, B. E. E., Rimmelzwaan, G. F. and Osterhaus, A. D. M. E. (2001). Infection of grey seals and harbour seals with influenza B virus. International Congress Series 1219, 225–231. doi: 10.1016/S0531-5131(01)00647-1.

Gabel, M., Theisen, S., Palm, H. W., Dähne, M. and Unger, P. (2021). Nematode Parasites in Baltic Sea Mammals, Grey Seal (Halichoerus grypus (Fabricius, 1791)) and Harbour Porpoise (Phocoena phocoena (L.)), from the German Coast. Acta Parasitologica 66, 26–33. doi: 10.1007/s11686-020-00246-7.

García de los Ríos y Loshuertos, A., Soler Laguía, M., Arencibia Espinosa, A., López Fernández, A., Covelo Figueiredo, P., Martínez Gomariz, F., Sánchez Collado, C., García Carrillo, N. and Ramírez Zarzosa, G. (2021). Comparative Anatomy of the Nasal Cavity in the Common Dolphin Delphinus delphis L., Striped Dolphin Stenella coeruleoalba M. and Pilot Whale Globicephala melas T.: A Developmental Study. Animals 11, 441. doi: 10.3390/ani11020441.

Garner, M. M., Lambourn, D. M., Jeffries, S. J., Hall, P. B., Rhyan, J. C., Ewalt, D. R., Polzin, L. M. and Cheville, N. F. (1997). Evidence of Brucella Infection in Parafilaroides Lungworms in a Pacific Harbor Seal (Phoca Vitulina Richardsi). Journal of Veterinary Diagnostic Investigation 9, 298–303. doi: 10.1177/104063879700900311.

Geraci, J. R. and St. Aubin, D. J. (1987). Effects of parasites on marine mammals. International Journal for Parasitology 17, 407–414. doi: 10.1016/0020-7519(87)90116-0.

Gerber, J. A., Roletto, J., Morgan, L. E., Smith, D. M. and Gage, L. J. (1993). Findings in pinnipeds stranded along the central and northern California coast, 1984–1990. Journal of Wildlife Diseases 29, 423–433. doi: 10.7589/0090-3558-29.3.423.

Gibson, D. I. and Harris, E. A. (1979). The helminth-parasites of cetaceans in the collection of the British Museum (Natural History). Investigations on Cetacea 10, 309–324.

Gibson, D. I., Harris, E. A., Bray, R. A., Jepson, P. D., Kuiken, T., Baker, J. R. and Simpson, V. R. (1998). A survey of the helminth parasites of cetaceans stranded on the coast of England and Wales during the period 1990-1994. Journal of Zoology 244, 563–574. doi: 10.1111/j.1469-7998.1998.tb00061.x.

Giorda, F., Ballardini, M., Di Guardo, G., Pintore, M. D., Grattarola, C., Iulini, B., Mignone, W., Goria, M., Serracca, L., Varello, K., Dondo, A., Acutis, P. L., Garibaldi, F., Scaglione, F. E., Gustinelli, A., Mazzariol, S., Di Francesco, C. E., Tittarelli, C., Casalone, C. and Pautasso, A. (2017). Postmortem findings in cetaceans found stranded in the pelagos sanctuary, Italy, 2007–14. Journal of Wildlife Diseases 53, 795. doi: 10.7589/2016-07-150.

Goldstein, T., Colegrove, K., Hanson, M. and Gulland, F. (2011). Isolation of a novel adenovirus from California sea lions Zalophus californianus. Diseases of Aquatic Organisms 94, 243–248. doi: 10.3354/dao02321.

Goodall, R., Galeazzi, A., Leatherwood, S., Miller, K., Cameron, I., Kastelein, R. and Sobral, A. (1988). Studies of Commerson’s dolphins, Cephalorhynchus commersonii, off Tierra del Fuego, 1976-1984, with a review of information on the species in the South Atlantic. Report of the International Whaling Commission 3–70.

Gosselin, J.-F. and Measures, L. N. (1997). Redescription of Filaroides (Parafilaroides) gymnurus(Railliet, 1899) (Nematoda: Metastrongyloidea), with comments on other species in pinnipeds. Canadian Journal of Zoology 75, 359–370. doi: 10.1139/z97-045.

Gosselin, J.-F., Measures, L. N. and Huot, J. (1998). Lungworm (Nematoda: Metastrongyloidea) infections in Canadian phocids. Canadian Journal of Fisheries and Aquatic Sciences 55, 825–834. doi: 10.1139/f97-306.

Greenwood, A. G. and Taylor, D. C. (1979). Odontocete parasites: Some new host records. Aquatic Mammals 7, 23–25.

Greig, D. J., Gulland, F. M. D. and Kreuder, C. (2005). A Decade of Live California Sea Lion (Zalophus californianus) Strandings Along the Central California Coast: Causes and Trends, 1991-2000. Aquatic Mammals 31, 11–22. doi: 10.1578/AM.31.1.2005.11.

Greig, D., Gulland, F., Smith, W., Conrad, P., Field, C., Fleetwood, M., Harvey, J., Ip, H., Jang, S., Packham, A., Wheeler, E. and Hall, A. (2014). Surveillance for zoonotic and selected pathogens in harbor seals Phoca vitulina from central California. Diseases of Aquatic Organisms 111, 93–106. doi: 10.3354/dao02762.

Groch, K. R., Santos-Neto, E. B., Díaz-Delgado, J., Ikeda, J. M. P., Carvalho, R. R., Oliveira, R. B., Guari, E. B., Bisi, T. L., Azevedo, A. F., Lailson-Brito, J. and Catão-Dias, J. L. (2018). Guiana Dolphin Unusual Mortality Event and Link to Cetacean Morbillivirus, Brazil. Emerging Infectious Diseases 24, 1349–1354. doi: 10.3201/eid2407.180139.

Groch, K. R., Díaz-Delgado, J., Santos-Neto, E. B., Ikeda, J. M. P., Carvalho, R. R., Oliveira, R. B., Guari, E. B., Flach, L., Sierra, E., Godinho, A. I., Fernández, A., Keid, L. B., Soares, R. M., Kanamura, C. T., Favero, C., Ferreira-Machado, E., Sacristán, C., Porter, B. F., Bisi, T. L., Azevedo, A. F., Lailson-Brito, J. and Catão-Dias, J. L. (2020a). The Pathology of Cetacean Morbillivirus Infection and Comorbidities in Guiana Dolphins During an Unusual Mortality Event (Brazil, 2017– 2018). Veterinary Pathology 57, 845–857. doi: 10.1177/0300985820954550.

Groch, K. R., Díaz-Delgado, J., Santos-Neto, E. B., Ikeda, J. M. P., Carvalho, R. R., Oliveira, R. B., Guari, E. B., Flach, L., Sierra, E., Godinho, A. I., Fernández, A., Keid, L. B., Soares, R. M., Kanamura, C. T., Favero, C., Ferreira-Machado, E., Sacristán, C., Porter, B. F., Bisi, T. L., Azevedo, A. F., Lailson-Brito, J. and Catão-Dias, J. L. (2020b). The Pathology of Cetacean Morbillivirus Infection and Comorbidities in Guiana Dolphins During an Unusual Mortality Event (Brazil, 2017– 2018). Veterinary Pathology 57, 845–857. doi: 10.1177/0300985820954550.

Gui, D., He, J., Zhang, X., Tu, Q., Chen, L., Feng, K., Liu, W., Mai, B. and Wu, Y. (2018). Potential association between exposure to legacy persistent organic pollutants and parasitic body burdens in Indo-Pacific finless porpoises from the Pearl River Estuary, China. Science of The Total Environment 643, 785–792. doi: 10.1016/j.scitotenv.2018.06.249.

Guimarães, J. P., Batista, R. L. G., Mariani, D. B. and Vergara-Parente, J. E. (2013). Ingestion of plastic debris by estuarine dolphin, Sotalia guianensis, off northeastern Brazil. Arquivos de Ciências do Mar 46, 107–122.

Guimarães, J. P., Febronio, A. M. B., Vergara-Parente, J. E. and Werneck, M. R. (2015). Lesions Associated with Halocercus brasiliensis Lins de Almeida, 1933 in the Lungs of Dolphins Stranded in the Northeast of Brazil. Journal of Parasitology 101, 248–251. doi: 10.1645/14-513.1.

Gulland, F. M. D., Beckmen, K., Burek, K., Lowenstine, L., Werner, L., Spraker, T., Dailey, M. and Harris, E. (1997a). Nematode (Otostrongylus circumlitus) infestation of northern elephant seals (Mirounga angustirostris) stranded along the central California coast. Marine Mammal Science 13, 446–458. doi: 10.1111/j.1748-7692.1997.tb00651.x.

Gulland, F. M. D., Beckmen, K., Burek, K., Lowenstine, L., Werner, L., Spraker, T., Dailey, M. and Harris, E. (1997b). NEMATODE (OTOSTRONGYLUS CIRCUMLITUS) INFESTATION OF NORTHERN ELEPHANT SEALS (MIROUNGA ANGUSTIROSTRIS) STRANDED ALONG THE CENTRAL CALIFORNIA COAST. Marine Mammal Science 13, 446–458. doi: 10.1111/j.1748-7692.1997.tb00651.x.

Gulland, F. M. D., Hall, A. J., Greig, D. J., Frame, E. R., Colegrove, K. M., Booth, R. K. N., Wasser, S. K. and Scott-Moncrieff, J. C. R. (2012). Evaluation of circulating eosinophil count and adrenal gland function in California sea lions naturally exposed to domoic acid. Journal of the American Veterinary Medical Association 241, 943–949. doi: 10.2460/javma.241.7.943.

Haddaway, N. R. and Watson, M. J. (2016). On the benefits of systematic reviews for wildlife parasitology. International Journal for Parasitology. Parasites and Wildlife 5, 184–191. doi: 10.1016/j.ijppaw.2016.05.002.

Harris, E. (1982). The helminth parasites of the Cetacea (or Parasitology with a porpoise).pp. R71–R71. Cambridge Univ Press.

Haulena, M. and Gulland, F. M. D. (2001). Use of medetomidine-zolazepam-tiletamine with and without atipamezole reversal to immobilize captive california sea lions. Journal of Wildlife Diseases 37, 566–573. doi: 10.7589/0090-3558-37.3.566.

Hermosilla, C., Silva, L. M. R., Prieto, R., Kleinertz, S., Taubert, A. and Silva, M. A. (2015). Endo- and ectoparasites of large whales (Cetartiodactyla: Balaenopteridae, Physeteridae): Overcoming difficulties in obtaining appropriate samples by non- and minimally-invasive methods. International Journal for Parasitology: Parasites and Wildlife 4, 414–420. doi: 10.1016/j.ijppaw.2015.11.002.

Hermosilla, C., Silva, L. M. R., Navarro, M. and Taubert, A. (2016). Anthropozoonotic Endoparasites in Free-Ranging “Urban” South American Sea Lions (*Otaria flavescens*). Journal of Veterinary Medicine 2016, 1–7. doi: 10.1155/2016/7507145.

Hernández-Orts, J. S., Hernández-Mena, D. I., Pantoja, C., Kuchta, R., García, N. A., Crespo, E. A. and Loizaga, R. (2021). A Visitor of Tropical Waters: First Record of a Clymene Dolphin (Stenella clymene) Off the Patagonian Coast of Argentina, With Comments on Diet and Metazoan Parasites. Frontiers in Marine Science 8, 658975. doi: 10.3389/fmars.2021.658975.

Herreman, J. K., McIntosh, A. D., Dziuba, R. K., Blundell, G. M., Ben-David, M. and Greiner, E. C. (2011). Parasites of harbor seals (Phoca vitulina) in Glacier Bay and Prince William Sound, Alaska. Marine Mammal Science 27, 247–253. doi: 10.1111/j.1748-7692.2009.00355.x.

Houde, M., Measures, L. N. and Huot, J. (2003). Lungworm (Pharurus pallasii: Metastrongyloidea: Pseudaliidae) infection in the endangered St. Lawrence beluga whale (Delphinapterus leucas). Canadian Journal of Zoology 81, 543–551.

Hsü, H. and Hoeppli, R. (1933). On some parasitic nematodes collected in Amoy. Peking Natural History Bulletin 8, 155–168.

Huertas, V. and Lagueux, C. J. (2016). First Recorded Mass Stranding of the Short-Finned Pilot Whale (Globicephala macrorhynchus) on the Caribbean Coast of Nicaragua. Aquatic Mammals 42, 27–34. doi: 10.1578/AM.42.1.2016.27.

Jabbar, A., Mohandas, N. and Gasser, R. B. (2014). Characterisation of the mitochondrial genome of Parafilaroides normani (lungworm) of Arctocephalus pusillus doriferus (Australian fur seal). Parasitology Research 113, 3049–3055. doi: 10.1007/s00436-014-3968-8.

Jacobus, K., Marigo, J., Gastal, S. B., Taniwaki, S. A., Ruoppolo, V., Catão-Dias, J. L. and Tseng, F. (2016). Identification of respiratory and gastrointestinal parasites of three species of pinnipeds Arctocephalus australis, Arctocephalus gazella, and Otaria flavescens in southern Brazil. Journal of Zoo and Wildlife Medicine 47, 132–140. doi: 10.1638/2015-0090.1.

Jauniaux, T., Petitjean, D., Brenez, C., Borrens, M., Brosens, L., Haelters, J., Tavernier, T. and Coignoul, F. (2002). Post-mortem Findings and Causes of Death of Harbour Porpoises (Phocoena phocoena) Stranded from 1990 to 2000 along the Coastlines of Belgium and Northern France. Journal of Comparative Pathology 126, 243–253. doi: 10.1053/jcpa.2001.0547.

Jepson, P. D., Kuiken, T., Bennett, P. M., Baker, J. R., Simpson, V. R. and Kennedy, S. (2000). Pulmonary pathology of harbour porpoises (Phocoena phocoena) stranded in England and Wales between 1990 and 1996. Veterinary Record 146, 721–728. doi: 10.1136/vr.146.25.721.

Jepson, P. D., Deaville, R., Barber, J. L., Aguilar, À., Borrell, A., Murphy, S., Barry, J., Brownlow, A., Barnett, J., Berrow, S., Cunningham, A. A., Davison, N. J., Ten Doeschate, M., Esteban, R., Ferreira, M., Foote, A. D., Genov, T., Giménez, J., Loveridge, J., Llavona, Á., Martin, V., Maxwell, D. L., Papachlimitzou, A., Penrose, R., Perkins, M. W., Smith, B., De Stephanis, R., Tregenza, N., Verborgh, P., Fernandez, A. and Law, R. J. (2016). PCB pollution continues to impact populations of orcas and other dolphins in European waters. Scientific Reports 6, 1–17. doi: 10.1038/srep18573.

Johnston, D. G. and Ridgway, S. H. (1969). Parasitism in some marine mammals. Journal of the American Veterinary Medical Association 155, 1064–1072.

Kastelein, R. A. and Lavaleije, M. S. S. (1992). Foreign Bodies in the stomach of female Habrour porpoises (Phocoena phocoena) from the North Sea. Aquatic Mammals 18, 40– 46.

Kaye, S., Johnson, S., Arnold, R. D., Nie, B., Davis, J. T., Gulland, F., Abou-Madi, N. and Fletcher, D. J. (2016). Pharamcokinetic study of oral e-aminocaproic acid in the northern elephant seal (Mirounga Angustirostris). Journal of Zoo and Wildlife Medicine 47, 438–446. doi: 10.1638/2015-0138.1.

Kaye, S., Johnson, S., Rios, C. and Fletcher, D. J. (2017). Plasmatic coagulation and fibrinolysis in healthy and Otostrongylus -affected Northern elephant seals (Mirounga angustirostris). Veterinary Clinical Pathology 46, 589–596. doi: 10.1111/vcp.12540.

Kelly, T. R., Greig, D., Colegrove, K. M., Lowenstine, L. J., Dailey, M., Gulland, F. M. and Haulena, M. (2005). Metastrongyloid Nematode (Otostrongylus circumlitus) Infection in a Stranded California Sea Lion (Zalophus californianus)— a New Host-parasite Association. Journal of Wildlife Diseases 41, 593–598. doi: 10.7589/0090-3558-41.3.593.

Kennedy, M. J. (1986). Metastrongyloidea) from the lungs of the ringed seal, Phoca hispida (Phocidae), from the Beaufort Sea, Canada. Canadian Journal of Zoology 64, 1864– 1868. doi: 10.1139/z86-278.

Kennedy, S., Smyth, J. A., Cush, P. F., Duignan, P., Platten, M., McCullough, S. J. and Allan, G. M. (1989). Histopathologic and Immunocytochemical Studies of Distemper in Seals. Veterinary Pathology 26, 97–103. doi: 10.1177/030098588902600201.

Kenyon, A. J. and Kenyon, B. J. (1977). Prevalence of Pharurus pallasii in the beluga whale (Delphinapterus leucas) of Churchill River Basin, Manitoba. Journal of Wildlife Diseases 13, 338–340. doi: 10.7589/0090-3558-13.4.338.

Kijewska, A. P., Jankowski, Z., Kuklik, I. and Rokicki, J. (2003). Pathological changes in the auditory organs of the harbor porpoise (Phocoena phocoena, L.) associated with Stenurus minor (Kuhn, 1829). Acta Parasitologica 48, 60–63. doi: 10.13140/RG.2.1.4781.7046.

Kontrimavichus, V. and Delyamure, S. (1979). Filaroides of domestic and wild animals. Fundamentals of nematology 29, 30–36.

Kroese, M. V., Beckers, L., Bisselink, Y. J. W. M., Brasseur, S., van Tulden, P. W., Koene, M. G. J., Roest, H. I. J., Ruuls, R. C., Backer, J. A., IJzer, J., van der Giessen, J. W. B. and Willemsen, P. T. J. (2018). Brucella pinnipedialis in grey seals (Halichoerus grypus) and harbour seals (Phoca vitulina) in the Netherlands. Journal of Wildlife Diseases 54, 439. doi: 10.7589/2017-05-097.

Kühn, J. (1829). Description d’un nouvelle espèce de strongyle trouveé dans le marsouin. Bulletin du Sciences Naturelles et Geologie 17, 150–153.

Kumazawa, H., Yona, R., Hirai, M. and Hasegawa, H. (2006). A Fatal Case of Bronchopneumonia Associated with Lungworm lnfection in a Bottle-nosed dolphin, tursiops truncatus (Cetacea: Dephinidae). Japanese Journal of Zoo and Wildlife Medicine 11, 31–34.

Kuramochi, T., Araki, J. and Machida, M. (1990). Pseudaliid nematodes from Dall’s porpoise, Phocoenoides dalli. Bulletin of the National Science Museum Series A (Zoology 97–103.

Kuramochi, T., Kikuchi, T., Okamura, H., Tatsukawa, T., Doi, H., Nakamura, K., Yamada, T., Koda, Y., Yoshida, Y. and Matsuura, M. (2000). Parasitic helminth and epizoit fauna of finless porpoise in the Inland Sea of Japan and the western North Pacific with a preliminary note on faunal difference by host’s local population. Memoirs of the National Science Museum, Tokyo 33, 83–95.

Kurochkin, Y. V. and Zablotsky, V. (1958). On the helminth fauna of the Caspian seal [In Russian]. Trudy Astrakhan—skogo Zapovednika 1993 337–343.

Kuwamura, M., Sawamoto, O., Yamate, J., Aoki, M., Ohnishi, Y. and Kotani, T. (2007). Pulmonary Vascular Proliferation and Lungworm (Stenurus ovatus) in a Bottlenose Dolphin (Tursiops turncatus). Journal of Veterinary Medical Science 69, 531–533. doi: 10.1292/jvms.69.531.

Kuzmina, T. A., Spraker, T. R., Kudlai, O., Lisitsyna, O. I., Zabludovskaja, S. O., Karbowiak, G., Fontaine, C. and Kuchta, R. (2018). Metazoan parasites of California sea lions (Zalophus californianus): A new data and review. International Journal for Parasitology: Parasites and Wildlife 7, 326–334. doi: 10.1016/j.ijppaw.2018.09.001.

Lambourn, D. M., Garner, M., Ewalt, D., Raverty, S., Sidor, I., Jeffries, S. J., Rhyan, J. and Gaydos, J. K. (2013). Brucella pinnipedialis infections in pacific harbour seals Phoca vitulina richardsi from Washington State, USA. Journal of Wildlife Diseases 49, 802– 815. doi: 10.7589/2012-05-137.

Lane, E. P., de Wet, M., Thompson, P., Siebert, U., Wohlsein, P. and Plön, S. (2014). A Systematic Health Assessment of Indian Ocean Bottlenose (Tursiops aduncus) and Indo-Pacific Humpback (Sousa plumbea) Dolphins Incidentally Caught in Shark Nets off the KwaZulu-Natal Coast, South Africa. PLoS ONE 9,. doi: 10.1371/journal.pone.0107038.

Lehnert, K., Raga, J. and Siebert, U. (2005). Macroparasites in stranded and bycaught harbour porpoises from German and Norwegian waters. Diseases of Aquatic Organisms 64, 265–269. doi: 10.3354/dao064265.

Lehnert, K., Raga, J. A. and Siebert, U. (2007). Parasites in harbour seals (Phoca vitulina) from the German Wadden Sea between two Phocine Distemper Virus epidemics. Helgoland Marine Research 61, 239–245. doi: 10.1007/s10152-007-0072-9.

Lehnert, K., von Samson-Himmelstjerna, G., Schaudien, D., Bleidorn, C., Wohlsein, P. and Siebert, U. (2010). Transmission of lungworms of harbour porpoises and harbour seals: Molecular tools determine potential vertebrate intermediate hosts. International Journal for Parasitology 40, 845–853. doi: 10.1016/j.ijpara.2009.12.008.

Lehnert, K., Seibel, H., Hasselmeier, I., Wohlsein, P., Iversen, M., Nielsen, N. H., Heide-Jørgensen, M. P., Prenger-Berninghoff, E. and Siebert, U. (2014). Increase in parasite burden and associated pathology in harbour porpoises (Phocoena phocoena) in West Greenland. Polar Biology 37, 321–331. doi: 10.1007/s00300-013-1433-2.

Lehnert, K., Randhawa, H. and Poulin, R. (2017). Metazoan parasites from odontocetes off New Zealand: new records. Parasitology Research 116, 2861–2868. doi: 10.1007/s00436-017-5573-0.

Lehnert, K., Poulin, R. and Presswell, B. (2019). Checklist of marine mammal parasites in New Zealand and Australian waters. Journal of Helminthology 93, 649–676. doi: 10.1017/S0022149X19000361.

Leidenberger, S. and Boström, S. (2009). Description of the lungworm Otostrongylus circumlitus (Railliet, 1899) de Bruyn, 1933 (Metastrongyloidea: Crenosomatidae) found in the heart of harbour seals from Sweden. Journal of Nematode Morphology and Systematics 12, 169–175.

Lipscomb, T. P., Kennedy, S., Moffett, D., Krafft, A., Klaunberg, B. A., Lichy, J. H., Regan, G. T., Worthy, G. A. J. and Taubenberger, J. K. (1996). Morbilliviral Epizootic in Bottlenose Dolphins of the Gulf of Mexico. Journal of Veterinary Diagnostic Investigation 8, 283–290. doi: 10.1177/104063879600800302.

Liu, H., Plancarte, M., Ball, E. E., Weiss, C. M., Gonzales-Viera, O., Holcomb, K., Ma, Z.-M., Allen, A. M., Reader, J. R., Duignan, P. J., Halaska, B., Khan, Z., Kriti, D., Dutta, J., van Bakel, H., Jackson, K., Pesavento, P. A., Boyce, W. M. and Coffey, L. L. (2021). Respiratory Tract Explant Infection Dynamics of Influenza A Virus in California Sea Lions, Northern Elephant Seals, and Rhesus Macaques. Journal of Virology 95,. doi: 10.1128/JVI.00403-21.

Lucas, Z., Daoust, P.-Y., Conboy, G. and Brimacombe, M. (2003). Health status of harp seals (Phoca groenlandica) and hooded seals (Cystophora cristata) on Sable Island, Nova Scotia, Canada, concurrent with their expanding range. Journal of Wildlife Diseases 39, 16–28. doi: 10.7589/0090-3558-39.1.16.

MacNeill, A. C., Neufeld, J. L. and Webster, W. A. (1975). Pulmonary nematodiasis in a narwhale. The Canadian Veterinary Journal = La Revue Veterinaire Canadienne 16, 53– 55.

Marigo, J., Ruoppolo, V., Rosas, F. C. W., Valente, A. L. S., Oliveira, M. R., Dias, R. A. and Catão-Dias, J. L. (2010). Helminths of Sotalia guianensis (Cetacea: Delphinidae) from the South and Southeastern Coasts of Brazil. Journal of Wildlife Diseases 46, 599–602. doi: 10.7589/0090-3558-46.2.599.

Mawson, P. M. (1953). Parasitic Nematoda collected by the Australian National Antarctic Research Expedition: Heard Island and Macquarie Island, 1948–1951. Parasitology 43, 291–297. doi: 10.1017/S0031182000018667.

Mazzariol, S., Marruchella, G., Di Guardo, G., Podesta, M., Olivieri, V., Colangelo, P., Kennedy, S., Castagnaro, M. and Cozzi, B. (2007). Post-mortem Findings in Cetacean Stranded along Italian Adriatic Sea coastline (2000–2006).

McColl, K. A. and Obendorf, D. L. (1982). Helminth parasites and associated pathology in stranded frasers dolphins, Lagenodelohis Hosei (Fraser, 1956). Aquatic Mammals 9, 30–34.

McFee, W. and Lipscomb, T. P. (2009). Major pathological findings and probable causes of mortality in bottlenose dolphins stranded in South Carolina from 1993 to 2006. Journal of Wildlife Diseases 45, 575–593. doi: 10.7589/0090-3558-45.3.575.

McKenzie, J. and Blair, D. (1983). Parasites from Hector’s dolphin (Cephalorhynchus hectori).pp. 126–127. Sir Publishing po box 399, Wellington, New zealand.

McKnight, C. A., Reynolds, T. L., Haulena, M., deLahunta, A. and Gulland, F. M. D. (2005). Congenital Hemicerebral Anomaly in a Stranded Pacific Harbor Seal (Phoca vitulina richardsi). Journal of Wildlife Diseases 41, 654–658. doi: 10.7589/0090-3558-41.3.654.

McManus, T., Wapstra, J., Guiler, E., Munday, B. and Obendorf, D. (1984). Cetacean strandings in Tasmania from February 1978 to May 1983. Papers and Proceedings of The Royal Society of Tasmania 118, 117–135. doi: 10.26749/rstpp.118.117.

Measures, L. N. (2001). Lungworms of Marine Mammals. In Parasitic Diseases of Wild Mammals (ed. Samuel, W. M., Pybus, M. J., and Kocan, A. A.), pp. 279–300. Iowa State University Press, Ames, Iowa, USA doi: 10.1002/9780470377000.ch10.

Measures, L. N. and Gosselin, J. (1994). Helminth parasites of ringed seal, Phoca hispida, from northern Quebec, Canada. Journal of the Helminthological Society of Washington 61, 240–244.

Measures, L. N., Béland, P., Martineau, D. and Guise, S. D. (1995). Helminths of an endangered population of belugas, Delphinapterus leucas, in the St. Lawrence estuary, Canada. Canadian Journal of Zoology 73, 1402–1409. doi: 10.1139/z95-165.

Melo, O. P., Ramos, R. M. A. and Di Beneditto, A. P. M. (2006). Helminths of the marine tucuxi, Sotalia fluviatilis (Gervais, 1853) (Cetacea: Delphinidae), in northern Rio de Janeiro State, Brazil. Brazilian Archives of Biology and Technology 49, 145–148. doi: 10.1590/S1516-89132006000100017.

Migaki, G., Van Dyke, D. and Hubbard, R. C. (1971). Some histopathological lesions caused by helminths in marine mammals. Journal of Wildlife Diseases 7, 281–289.

Mignucci-Giannoni, A. A., Rodríguez-López, M. A., Perez-Zayas, J. J., Montoya-Ospina, R. A. and Williams, E. H. J. (1998a). First record of the melonhead whale for Puerto Rico. Mammalia 62, 452–457.

Mignucci-Giannoni, A. A., Hoberg, E. P., Siegel-Causey, D. and Williams, E. H. (1998b). Metazoan Parasites and Other Symbionts of Cetaceans in the Caribbean. The Journal of Parasitology 84, 939. doi: 10.2307/3284625.

Morales Vela, B. and Olvera Gómez, L. D. (1993). Varamiento de calderones Globicephala macrorhynchus (Cetacea: Delphinidae) en la Isla de Cozumel, Quintana Roo, México. Anales del Instituto de Biología serie Zoología 64,.

Morell, M., Lehnert, K., IJsseldijk, L., Raverty, S., Wohlsein, P., Gröne, A., André, M., Siebert, U. and Shadwick, R. (2017). Parasites in the inner ear of harbour porpoise: cases from the North and Baltic Seas. Diseases of Aquatic Organisms 127, 57–63. doi: 10.3354/dao03178.

Moser, M. and Rhinehart, H. (1993). The lungworm, Halocercus spp. (Nematoda: Pseudaliidae) in cetaceans from California. Journal of Wildlife Diseases 29, 507–508. doi: 10.7589/0090-3558-29.3.507.

Nicholson, A. and Fanning, J. (1981). Parasites and associated pathology of the respiratory tract of the Australian sea lion: Neophoca cinerea.pp. 178–181.

Niebuhr, C. N., Siers, S. R., Leinbach, I. L., Kaluna, L. M. and Jarvi, S. I. (2021). Variation in *Angiostrongylus cantonensis* infection in definitive and intermediate hosts in Hawaii, a global hotspot of rat lungworm disease. Parasitology 148, 133–142. doi: 10.1017/S003118202000164X.

Nunn, C. L., Altizer, S., Jones, K. E. and Sechrest, W. (2003). Comparative Tests of Parasite Species Richness in Primates. The American Naturalist 162, 597–614. doi: 10.1086/378721.

Oliveira, J. B., Morales, J. A., González-Barrientos, R. C., Hernández-Gamboa, J. and Hernández-Mora, G. (2011). Parasites of cetaceans stranded on the Pacific coast of Costa Rica. Veterinary Parasitology 182, 319–328. doi: 10.1016/j.vetpar.2011.05.014.

Onderka, D. K. (1989). Prevance and pathology of nematode infections in the lungs of ringed seals (Phoca hispida) of the Western Arctic of Canada. Journal of Wildlife Diseases 25, 218–224. doi: 10.7589/0090-3558-25.2.218.

Osinga, N., Kappe, A. L., Brakefield, P. M., Udo de Haes, H. A. and Elson-Riggins, J. G. (2015). Comparative biology of common and grey seals along the Dutch coast: stranding, disease, rehabilitation and conservation. Chapter 8: Observations regarding transmission of seal nematodes in common seals, Phoca vitulina vitulina from the Wadden Sea.

Page, M. J., McKenzie, J. E., Bossuyt, P. M., Boutron, I., Hoffmann, T. C., Mulrow, C. D., Shamseer, L., Tetzlaff, J. M., Akl, E. A., Brennan, S. E., Chou, R., Glanville, J., Grimshaw, J. M., Hróbjartsson, A., Lalu, M. M., Li, T., Loder, E. W., Mayo-Wilson, E., McDonald, S., McGuinness, L. A., Stewart, L. A., Thomas, J., Tricco, A. C., Welch, V. A., Whiting, P. and Moher, D. (2021). The PRISMA 2020 statement: An updated guideline for reporting systematic reviews. PLOS Medicine 18, e1003583. doi: 10.1371/journal.pmed.1003583.

Parsons, E. C. M. and Jefferson, T. A. (2000). Post-mortem investigations on stranded dolphins and porpoises from Hong Kong waters. Journal of Wildlife Diseases 36, 342–356. doi: 10.7589/0090-3558-36.2.342.

Parsons, E. C. M., Overstreet, R. M. and Jefferson, T. A. (2001). Parasites from Indo-Pacific hump-backed dolphins (Sousa chinensis) and finless porpoises (Neophocaena phocaenoides) stranded in Hong Kong. Veterinary Record 148, 776–780. doi: 10.1136/vr.148.25.776.

Pedersen, A. B. and Fenton, A. (2015). The role of antiparasite treatment experiments in assessing the impact of parasites on wildlife. Trends in Parasitology 31, 200–211. doi: 10.1016/j.pt.2015.02.004.

Pekmezci, G. Z., Yardimci, B., Gürler, A. T., Bölükbaş, C. S., Açici, M. and Umur, Ş. (2013). Survey on the Presence of Nematodes and Associated with Pathology in Marine Mammals from Turkish Waters. Kafkas Universitesi Veteriner Fakultesi Dergisi 19, 1035–1038. doi: 10.9775/kvfd.2013.9409.

Perrin, W. F., Mitchell, E. D., Mead, J. G., Caldwell, D. K., Caldwell, M. C., Bree, P. J. H. and Dawbin, W. H. (1987). Revision of the spotted dolphins, Stenella spp. Marine Mammal Science 3, 99–170. doi: 10.1111/j.1748-7692.1987.tb00158.x.

Perrin, W. F., Caldwell, D. K. and Caldwell, M. (1994). Atlantic spotted dolphin Stenella frontalis (G. Cuvier, 1829). Handbook of marine mammals 5, 173–190.

Petter, A. J. and Pilleri, G. (1982). Pharurus asiaeorientalis new species, metastrongylid nematode, parasite of Neophocaena asiaeorientalis (Phocoenidae, Cetacea). Investigations on Cetacea 13, 141–148.

Piché, C., Measures, L., Bédard, C. and Lair, S. (2010). Bronchoalveolar lavage and pulmonary histopathology in harp seals (Phoca groenlandica) experimentally infected with Otostrongylus circumlitis. Journal of Wildlife Diseases 46, 409–421. doi: 10.7589/0090-3558-46.2.409.

Pool, R., Chandradeva, N., Gkafas, G., Raga, J. A., Fernández, M. and Aznar, F. J. (2020a). Transmission and Predictors of Burden of Lungworms of the Striped Dolphin (Stenella coeruleoalba) in the Western Mediterranean. Journal of Wildlife Diseases 56, 186. doi: 10.7589/2018-10-242.

Pool, R., Fernández, M., Chandradeva, N., Raga, J. A. and Aznar, F. J. (2020b). The taxonomic status of Skrjabinalius guevarai Gallego & Selva, 1979 (Nematoda: Pseudaliidae) and the synonymy of Skrjabinalius Delyamure, 1942 and Halocercus Baylis & Daubney, 1925. Systematic Parasitology 97, 389–401. doi: 10.1007/s11230-020-09921-9.

Pool, R., Romero-Rubira, C., Raga, J. A., Fernández, M. and Aznar, F. J. (2021). Determinants of lungworm specificity in five cetacean species in the western Mediterranean. Parasites & Vectors 14, 196. doi: 10.1186/s13071-021-04629-1.

Poulin, R. (2011). Evolutionary Ecology of Parasites: (Second *Edition*). Princeton University Press doi: doi:10.1515/9781400840809.

Prahl, S., Ketten, D. R. and Siebert, U. (2008). Examinations of ears in Harbour porpoises Phocoena phocoena from the north and baltic seas. Bioacoustics 17, 85–87. doi: 10.1080/09524622.2008.9753775.

Prenger-Berninghoff, E., Siebert, U., Stede, M., König, A., Weiß, R. and Baljer, G. (2008). Incidence of Brucella species in marine mammals of the German North Sea. Diseases of Aquatic Organisms 81, 65–71. doi: 10.3354/dao01920.

Raga, J. A. (1994). Parasitismus bei den Cetacea. In Robineau D, Duguy R, Klima M *(eds.).* Handbuch der Säugetiere Europas, pp. 132–179.

Raga, J. and Balbuena, J. (1987). Algunas características zoogeográficas de los helmintos de los cetáceos en el Mediterráneo con especial referencia a la helmintofauna del delfín listado. Mamíferos y helmintos. Barcelona: Ketres Editora 195, 201.

Reckendorf, A., Ludes-Wehrmeister, E., Wohlsein, P., Tiedemann, R., Siebert, U. and Lehnert, K. (2018). First record of Halocercus sp. (Pseudaliidae) lungworm infections in two stranded neonatal orcas (Orcinus orca). Parasitology 145, 1553–1557. doi: 10.1017/S0031182018000586.

Reckendorf, A., Everaarts, E., Bunskoek, P., Haulena, M., Springer, A., Lehnert, K., Lakemeyer, J., Siebert, U. and Strube, C. (2021). Lungworm infections in harbour porpoises (Phocoena phocoena) in the German Wadden Sea between 2006 and 2018, and serodiagnostic tests. International Journal for Parasitology: Parasites and Wildlife 14, 53–61. doi: 10.1016/j.ijppaw.2021.01.001.

Regassa, A., Toyeb, M., Abebe, R., Megersa, B., Mekibib, B., Mekuria, S., Debela, E. and Abunna, F. (2010). Lungworm infection in small ruminants: Prevalence and associated risk factors in Dessie and Kombolcha districts, northeastern Ethiopia. Veterinary Parasitology 169, 144–148. doi: 10.1016/j.vetpar.2009.12.010.

Reisfeld, L., Sacristán, C., Sánchez-Sarmiento, A. M., Costa-Silva, S., Díaz-Delgado, J., Groch, K. R., Marigo, J., Ewbank, A. C., Favero, C. M., Guerra, J. M., Réssio, R. A., Cremer, M. J., Esperón, F. and Catão-Dias, J. L. (2019). Fatal pulmonary parafilaroidiasis in a free-ranging subantarctic fur seal (Arctocephalus tropicalis) coinfected with two gammaherpesviruses and Sarcocystis sp. Revista Brasileira de Parasitologia Veterinária 28, 499–503. doi: 10.1590/s1984-29612019029.

Reyes, J. C. and van Waerebeek, K. (1995). Aspects of the biology of Burmeister’s porpoise from Peru. Report of the International Whaling Commission 349–364.

Rhyan, J., Garner, M., Spraker, T., Lambourn, D. and Cheville, N. (2018a). Brucella pinnipedialis in lungworms Parafilaroides sp. and Pacific harbor seals Phoca vitulina richardsi: proposed pathogenesis. Diseases of Aquatic Organisms 131, 87–94. doi: 10.3354/dao03291.

Rhyan, J., Garner, M., Spraker, T., Lambourn, D. and Cheville, N. (2018b). Brucella pinnipedialis in lungworms Parafilaroides sp. and Pacific harbor seals Phoca vitulina richardsi: proposed pathogenesis. Diseases of Aquatic Organisms 131, 87–94. doi: 10.3354/dao03291.

Rodrigues, T., Díaz-Delgado, J., Catão-Dias, J., da Luz Carvalho, J. and Marmontel, M. (2018). Retrospective pathological survey of pulmonary disease in free-ranging Amazon river dolphin Inia geoffrensis and tucuxi Sotalia fluviatilis. Diseases of Aquatic Organisms 131, 1–11. doi: 10.3354/dao03280.

Rogan, E., Baker, J. R., Jepson, P. D., Berrow, S. and Kiely, O. (1997). A mass stranding of white-sided dolphins (Lagenorhynchus acutus) in Ireland: biological and pathological studies. Journal of Zoology 242, 217–227. doi: 10.1111/j.1469-7998.1997.tb05798.x.

Rogan, E., Penrose, R., Gassner, I., Mackey, M. J. and Clayton, P. (2001). Marine Mammal Strandings: A Collaborative Study for the Irish Sea. The Marine Institute.

Rosas, F. C. W., Monteiro-Filho, E. L. A., Marigo, J., Santos, R. A., Andrade, A. L. V., Rautenberg, M., Olivcira, M. R. and Bordignon, M. O. (2002). The striped dolphin, Stenella coeruleoalba (Cetacea: Delphinidae), on the coast of São Paulo State, southeastern Brazil. Aquatic Mammals 28, 60–66.

Ross, G. J. B. and Bass, A. J. (1971). Shark attack on an ailing dolphin, Stenella Coeruleoalba (Meyen). South African Journal of Science.

Santos, C., Rohde, K., Ramos, R., Di Beneditto, A. and Capistrano, L. (1996). Helminths of cetaceans on the Southeastern coast of Brazil. Journal of the Helminthological Society of Washington 63, 149–152.

Savage, K. N., Burek-Huntington, K., Wright, S. K., Bryan, A. L., Sheffield, G., Webber, M., Stimmelmayr, R., Tuomi, P., Delaney, M. A. and Walker, W. (2021). Stejneger’s beaked whale strandings in Alaska, 1995–2020. Marine Mammal Science 37, 843–869. doi: 10.1111/mms.12780.

Schick, L., IJsseldijk, L. L., Grilo, M. L., Lakemeyer, J., Lehnert, K., Wohlsein, P., Ewers, C., Prenger-Berninghoff, E., Baumgärtner, W., Gröne, A., Kik, M. J. L. and Siebert, U. (2020). Pathological Findings in White-Beaked Dolphins (Lagenorhynchus albirostris) and Atlantic White-Sided Dolphins (Lagenorhynchus acutus) From the South-Eastern North Sea. Frontiers in Veterinary Science 7, 262. doi: 10.3389/fvets.2020.00262.

Schmid-Hempel, P. (2021). Virulence evolution. In Evolutionary Parasitology, pp. 353–388. Oxford University Press doi: 10.1093/oso/9780198832140.003.0013.

Schumacher, U., Horny, H., Heidemann, G., Schultz, W. and Welsch, U. (1990). Histopathological findings in harbour seals (Phoca vitulina) found dead on the german north sea coast. Journal of Comparative Pathology 102, 299–309. doi: 10.1016/S0021-9975(08)80019-9.

Seguel, M. and Gottdenker, N. (2017). The diversity and impact of hookworm infections in wildlife. International Journal for Parasitology: Parasites and Wildlife 6, 177–194. doi: 10.1016/j.ijppaw.2017.03.007.

Seguel, M., Nadler, S., Field, C. and Duignan, P. (2018a). Vasculitis and Thrombosis due to the Sea Lion Lungworm, *Parafilaroides decorus*, in a Guadalupe Fur Seal (*Arctocephalus philippii townsendi*). Journal of Wildlife Diseases 54, 638–641. doi: 10.7589/2017-12-291.

Seguel, M., Nadler, S., Field, C. and Duignan, P. (2018b). Vasculitis and Thrombosis due to the Sea Lion Lungworm, *Parafilaroides decorus*, in a Guadalupe Fur Seal (*Arctocephalus philippii townsendi*). Journal of Wildlife Diseases 54, 638–641. doi: 10.7589/2017-12-291.

Seguel, M., George, R. C., Maboni, G., Sanchez, S., Page-Karjian, A., Wirth, E., McFee, W. and Gottdenker, N. L. (2020a). Pathologic findings and causes of death in bottlenose dolphins Tursiops truncatus stranded along the Georgia coast, USA (2007−2013). Diseases of Aquatic Organisms 141, 25–38. doi: 10.3354/dao03509.

Seguel, M., George, R., Maboni, G., Sanchez, S., Page-Karjian, A., Wirth, E., McFee, W. and Gottdenker, N. (2020b). Pathologic findings and causes of death in bottlenose dolphins Tursiops truncatus stranded along the Georgia coast, USA (2007-2013). Diseases of Aquatic Organisms 141, 25–38. doi: 10.3354/dao03509.

Seguel, M., Budischak, S. A., Jolles, A. E. and Ezenwa, V. O. (2023). Helminth-associated changes in host immune phenotype connect top-down and bottom-up interactions during co-infection. Functional Ecology 37, 860–872. doi: 10.1111/1365-2435.14237.

Seibel, H., Beineke, A. and Siebert, U. (2010). Mycotic Otitis Media in a Harbour Porpoise (Phocoena phocoena). Journal of Comparative Pathology 143, 294–296. doi: 10.1016/j.jcpa.2010.03.002.

Sheldon, J. D., Johnson, S. P., Hernandez, J. A., Cray, C. and Stacy, N. I. (2017). Acute-phase responses in healthy, malnourished, and Otostrongylus-infected juvenile northern elephant seals (mirounga angustirostris). Journal of Zoo and Wildlife Medicine 48, 767– 775. doi: 10.1638/2016-0267.1.

Sheldon, J. D., Hernandez, J. A., Johnson, S. P., Field, C., Kaye, S. and Stacy, N. I. (2019). Diagnostic Performance of Clinicopathological Analytes in Otostrongylus circumlitis-Infected Rehabilitating Juvenile Northern Elephant Seals (Mirounga angustirostris). Frontiers in Veterinary Science 6, 134. doi: 10.3389/fvets.2019.00134.

Shiozaki, A. and Amano, M. (2017). Population- and growth-related differences in helminthic fauna of finless porpoises (Neophocaena asiaeorientalis) in five Japanese populations. Journal of Veterinary Medical Science 79, 534–541. doi: 10.1292/jvms.16-0421.

Siebert, U., Wünschmann, A., Weiss, R., Frank, H., Benke, H. and Frese, K. (2001). Post-mortem Findings in Harbour Porpoises (Phocoena phocoena) from the German North and Baltic Seas. Journal of Comparative Pathology 124, 102–114. doi: 10.1053/jcpa.2000.0436.

Siebert, U., Tolley, K., Víkingsson, G. A., Ólafsdottir, D., Lehnert, K., Weiss, R. and Baumgärtner, W. (2006). Pathological Findings in Harbour Porpoises (Phocoena phocoena) from Norwegian and Icelandic Waters. Journal of Comparative Pathology 134, 134–142. doi: 10.1016/j.jcpa.2005.09.002.

Siebert, U., Jepson, P. D. and Wohlsein, P. (2013). First indication of gas embolism in a harbour porpoise (Phocoena phocoena) from German waters. European Journal of Wildlife Research 59, 441–444. doi: 10.1007/s10344-013-0700-4.

Siebert, U., Pawliczka, I., Benke, H., von Vietinghoff, V., Wolf, P., Pilāts, V., Kesselring, T., Lehnert, K., Prenger-Berninghoff, E., Galatius, A., Anker Kyhn, L., Teilmann, J., Hansen, M. S., Sonne, C. and Wohlsein, P. (2020). Health assessment of harbour porpoises (PHOCOENA PHOCOENA) from Baltic area of Denmark, Germany, Poland and Latvia. Environment International 143, 105904. doi: 10.1016/j.envint.2020.105904.

Sierra, E., Zucca, D., Arbelo, M., García-Álvarez, N., Andrada, M., Déniz, S. and Fernández, A. (2014). Fatal Systemic Morbillivirus Infection in Bottlenose Dolphin, Canary Islands, Spain. Emerging Infectious Diseases 20, 269–271. doi: 10.3201/eid2002.131463.

Smith, F. R. and Threlfall, W. (1973). Helminths of Some Mammals from Newfoundland. American Midland Naturalist 90, 215. doi: 10.2307/2424284.

Stephens, N., Duignan, P. J., Wang, J., Bingham, J., Finn, H., Bejder, L., Patterson, A. P. and Holyoake, C. (2014). Cetacean Morbillivirus in Coastal Indo-Pacific Bottlenose Dolphins, Western Australia. Emerging Infectious Diseases 20, 672–676. doi: 10.3201/eid2004.131714.

Stijnen, T., Hamza, T. H. and Özdemir, P. (2010). Random effects meta-analysis of event outcome in the framework of the generalized linear mixed model with applications in sparse data. Statistics in Medicine 29, 3046–3067. doi: 10.1002/sim.4040.

Stockin, K. A., Duignan, P. J., Roe, W. A., Meynier, L., Alley, M. and Fettermann, T. (2009). Causes of mortality in stranded Common Dolphin (Delphinus sp.) from New Zealand waters between 1998 and 2008. Pacific Conservation Biology 15, 217. doi: 10.1071/PC090217.

Stroud, R. K. (1978). Parasites and associated pathology observed in pinnipeds stranded along the Oregon coast. Journal of Wildlife Diseases 14, 292–298. doi: 10.7589/0090-3558-14.3.292.

Stroud, R. K. and Roffe, T. J. (1979). Causes of death in marine mammals stranded along the oregon coast. Journal of Wildlife Diseases 15, 91–97. doi: 10.7589/0090-3558-15.1.91.

Suvorova, I. V. and Prokushina, K. S. (2021). Pulmonary nematodiasis of the Baikal seal (Pusa siberica). In Theory and practice of parasitic disease control: Collection of Scientific Articles adapted from the International Scientific Conference., pp. 509–514.

Sweeney, J. C. and Gilmartin, W. G. (1974). Survey of diseases in free-living California sea lions. Journal of Wildlife Diseases 10, 370–376. doi: 10.7589/0090-3558-10.4.370.

Szefer, P., Rokicki, J., Frelek, K., Skora, K. and Malinga, M. (1998). Bioaccumulation of selected trace elements in lung nematodes, Pseudalius inflexus, of harbor porpoise (Phocoena phocoena) in a Polish zone of the Baltic Sea. The Science of the Total Environment 220, 19–24.

Tao, J.-Y. (1983). A new species and a new chinese record of nematodes from porpoise neophocaenaphocaenoides. Acta Zootaxonomica Sinica.

Taylor, M. A., Coop, R. L. and Wall, R. L. eds. (2015). Veterinary Helminthology. In Veterinary Parasitology, pp. 1–109. John Wiley & Sons, Inc., Hoboken, NJ, USA doi: 10.1002/9781119073680.ch1.

Terracciano, G., Fichi, G., Comentale, A., Ricci, E., Mancusi, C. and Perrucci, S. (2020). Dolphins Stranded along the Tuscan Coastline (Central Italy) of the “Pelagos Sanctuary”: A Parasitological Investigation. Pathogens 9, 612. doi: 10.3390/pathogens9080612.

Tomilin, A. (1967). Mammals of the USSR and adjacent countries, Vol. 9, Cetacea. Israel program for scientific translations, Jerusalem 71,.

Tomo, I., Kemper, C. M. and Lavery, T. J. (2010). Eighteen-year study of south Australian dolphins shows variation in lung nematodes by season, year, age class, and location. Journal of Wildlife Diseases 46, 488–498. doi: 10.7589/0090-3558-46.2.488.

Torres, P., Cortes, P., Oporto, J. A., Brieva, L. and Silva, R. (1994). The occurrence of stenurus australis Tantalean and Sarmiento, 1991(Nematoda: Mestratongyloidea)in the Porpoise Phocoena spinipinnis (Burmeister,1865) on the Southern Coast of Chile. Memórias do Instituto Oswaldo Cruz 89, 141–143. doi: 10.1590/S0074-02761994000200004.

Ulrich, S. A., Lehnert, K., Siebert, U. and Strube, C. (2015). A recombinant antigen-based enzyme-linked immunosorbent assay (ELISA) for lungworm detection in seals. Parasites & Vectors 8, 443. doi: 10.1186/s13071-015-1054-4.

Ulrich, S. A., Lehnert, K., Rubio-Garcia, A., Sanchez-Contreras, G. J., Strube, C. and Siebert, U. (2016). Lungworm seroprevalence in free-ranging harbour seals and molecular characterisation of marine mammal MSP. International Journal for Parasitology: Parasites and Wildlife 5, 48–55. doi: 10.1016/j.ijppaw.2016.02.001.

Valderrama Vasquez, C. A., Macgregor, S. K., Rowcliffe, J. M. and Jepson, P. D. (2008). Occurrence of a monophasic strain of Salmonella group B isolated from cetaceans in England and Wales between 1990 and 2002. Environmental Microbiology 10, 2462–2468. doi: 10.1111/j.1462-2920.2008.01651.x.

van Elk, C. E., van de Bildt, M. W. G., van Run, P. R. W. A., Bunskoek, P., Meerbeek, J., Foster, G., Osterhaus, A. D. M. E. and Kuiken, T. (2019). Clinical, pathological, and laboratory diagnoses of diseases of harbour porpoises (Phocoena phocoena), live stranded on the Dutch and adjacent coasts from 2003 to 2016. Veterinary Research 50, 88. doi: 10.1186/s13567-019-0706-3.

van Wijngaarden, M. F. A., Geut, M. I. M., Vernooij, J. C. M., IJsseldijk, L. L. and Tobias, T. J. (2021). Determinants of mortality of juvenile harbour seals (Phoca vitulina) infected with lungworm submitted to a Dutch seal rehabilitation centre. International Journal for Parasitology: Parasites and Wildlife 14, 1–6. doi: 10.1016/j.ijppaw.2020.12.002.

Vargas-Castro, I., Crespo-Picazo, J. L., Rivera-Arroyo, B., Sánchez, R., Marco-Cabedo, V., Jiménez-Martínez, M. Á., Fayos, M., Serdio, Á., García-Párraga, D. and Sánchez-Vizcaíno, J. M. (2020). Alpha- and gammaherpesviruses in stranded striped dolphins (Stenella coeruleoalba) from Spain: first molecular detection of gammaherpesvirus infection in central nervous system of odontocetes. BMC Veterinary Research 16, 288. doi: 10.1186/s12917-020-02511-3.

Vercruysse, J., Salomez, A., Ulloa, A., Osterhaus, A., Kuiken, T. and Alvinerie, M. (2003). Efficacy of ivermectin and moxidectin against Otostrongylus circumlitus and Parafilaroides gymnurus in harbour seals (Phoca vitulina). Veterinary Record 152, 130–134. doi: 10.1136/vr.152.5.130.

Veryeri, N. G. (2012). Postmortem examinations of stranded dolphins found on the Black Sea coast near Ordu, Turkey (Mammalia: Cetacea). Zoology in the Middle East 55, 129–132. doi: 10.1080/09397140.2012.10648928.

von Waerebeek, K., Reyes, J. C. and Shigueto, J. A. (1993). Helminth parasites and phoronts of dusky dolphins Lagenorhynchus obscurus (Gray, 1828) from Peru. Aquatic Mammals 19, 159–169.

Walden, H. D. S., Grijalva, C. J., Páez-Rosas, D. and Hernandez, J. A. (2018). Intestinal Parasites in Galapagos Sea Lions (Zalophus wollebaeki) Sivertsen, 1953 on San Cristóbal Island, Galapagos, Ecuador. Journal of Parasitology 104, 718–721. doi: 10.1645/17-187.

Walden, H. S., Bryan, A. L., McIntosh, A., Tuomi, P., Hoover-Miller, A., Stimmelmayr, R. and Quakenbush, L. (2020). Helminth fauna of ice seals in the Alaskan Bering and Chukchi seas, 2006–15. Journal of Wildlife Diseases 56,. doi: 10.7589/2019-09-228.

Walker, W. A. (1975). Review of the Live-Capture Fishery for Smaller Cetaceans Taken in Southern California Waters for Public Display, 1966–73. Journal of the Fisheries Research Board of Canada 32, 1197–1211. doi: 10.1139/f75-139.

Wan, X., Zheng, J., Li, W., Zeng, X., Yang, J., Hao, Y. and Wang, D. (2017). Parasitic infections in the East Asian finless porpoise Neophocaena asiaeorientalis sunameri living off the Chinese Yellow/Bohai Sea coast. Diseases of Aquatic Organisms 125, 63–71. doi: 10.3354/dao03131.

Wazura, K. W., Strong, J. T., Glenn, C. L. and Bush, A. O. (1986). Helminths of the Beluga Whale (Delphinapterus leucas) from the Mackenzie River Delta, Northwest Territories. Journal of Wildlife Diseases 22, 440–442. doi: 10.7589/0090-3558-22.3.440.

Webster, W., Neufeld, J. and MacNeill, A. (1973). Halocercus monoceris sp. n. (Nematoda: Metastrongyloidea) from the Narwhal, Monodon monoceros. Proceedings of the Helminthological Society of Washington 40, 255–258.

Williams, K. M., Fessler, M. K., Bloomfield, R. A., Sandke, W. D., Malekshahi, C. R., Keroack, C. D., Duignan, P. J., Torquato, S. D. and Williams, S. A. (2020). A novel quantitative real-time PCR diagnostic assay for fecal and nasal swab detection of an otariid lungworm, Parafilaroides decorus. International Journal for Parasitology: Parasites and Wildlife 12, 85–92. doi: 10.1016/j.ijppaw.2020.04.012.

Wohlsein, P., Seibel, H., Beineke, A., Baumgärtner, W. and Siebert, U. (2019a). Morphological and Pathological Findings in the Middle and Inner Ears of Harbour Porpoises (Phocoena phocoena). Journal of Comparative Pathology 172, 93–106. doi: 10.1016/j.jcpa.2019.09.005.

Wohlsein, P., Seibel, H., Beineke, A., Baumgärtner, W. and Siebert, U. (2019b). Morphological and Pathological Findings in the Middle and Inner Ears of Harbour Porpoises (Phocoena phocoena). Journal of Comparative Pathology 172, 93–106. doi: 10.1016/j.jcpa.2019.09.005.

Woodard, J. C., Zam, S. G., Caldwell, D. K. and Caldwell, M. C. (1969). Some Parasitic Diseases of Dolphins. Pathologia veterinaria 6, 257–272. doi: 10.1177/030098586900600307.

WoRMS Editorial Board (2022). World Register of Marine Species. doi: 10.14284/170.

Wu, H. W. (1929). On Halocercus pingi n. sp. a Lung-Worm from the Porpoise, Neomeris phocoenoides. The Journal of Parasitology 15, 276. doi: 10.2307/3271983.

Wunschimann, A., Frese, K., Müiller, G., Baumgärtner, W., Siebert, U., Weiss, R., Lockyer, C. and Heide-Jørgensen, M. P. (2001). Evidence of infectious diseases in harbour porpoises (Phocoena phocoena) hunted in the waters of Greenland and by-caught in the German North Sea and Baltic Sea. Veterinary Record 148, 715–720. doi: 10.1136/vr.148.23.715.

Wünschmann, A., Siebert, U. and Weiss, R. (1999). Rhizopusmycosis in a Harbor Porpoise from the Baltic Sea. Journal of Wildlife Diseases 35, 569–573. doi: 10.7589/0090-3558-35.3.569.

Yamaguti, S. (1951). Studies on the helminth fauna of Japan. Part 46. Nematodes of marine mammals. Arbeiten aus der Medizinischen Fakultat Okayama 7, 295–306.

Yu, J., Sun, Y. and Xia, Z. (2009). The Rescue, Rehabilitation, and Release of a Stranded Finless Porpoise (Neophocaena phocaenoides sunameri) at Bohai Bay of China. Aquatic Mammals 35, 220–225. doi: 10.1578/AM.35.2.2009.220.

Zafra, R., Jaber, J. R., Pérez, J., de la Fuente, J., Arbelo, M., Andrada, M. and Fernández, A. (2015a). Immunohistochemical characterisation of parasitic pneumonias of dolphins stranded in the Canary Islands. Research in Veterinary Science 100, 207–212. doi: 10.1016/j.rvsc.2015.03.021.

Zafra, R., Jaber, J. R., Pérez, J., de la Fuente, J., Arbelo, M., Andrada, M. and Fernández, A. (2015b). Immunohistochemical characterisation of parasitic pneumonias of dolphins stranded in the Canary Islands. Research in Veterinary Science 100, 207–212. doi: 10.1016/j.rvsc.2015.03.021.

Zam, S. G., Caldwell, D. K. and Caldwell, M. C. (1971). Some endoparasites from small odontocete cetaceans collected in Florida and Georgia. Cetology 2, 1–11.

Zylber, M. I., Failla, G. and Le Bas, A. (2002). Stenurus globicephalae Baylis et Daubney, 1925 (Nematoda: Pseudaliidae) from a False Killer Whale, Pseudorca crassidens (Cetacea: Delphinidae), Stranded on the Coast of Uruguay. Memórias do Instituto Oswaldo Cruz 97, 221–225. doi: 10.1590/S0074-02762002000200015.

