## Supplementary Figures for "A systematic review of the diversity and virulence correlates of metastrongyle lungworms in marine mammals"

Title:

Fischbach, J.R. & Seguel, M.

Supplementary Figures:

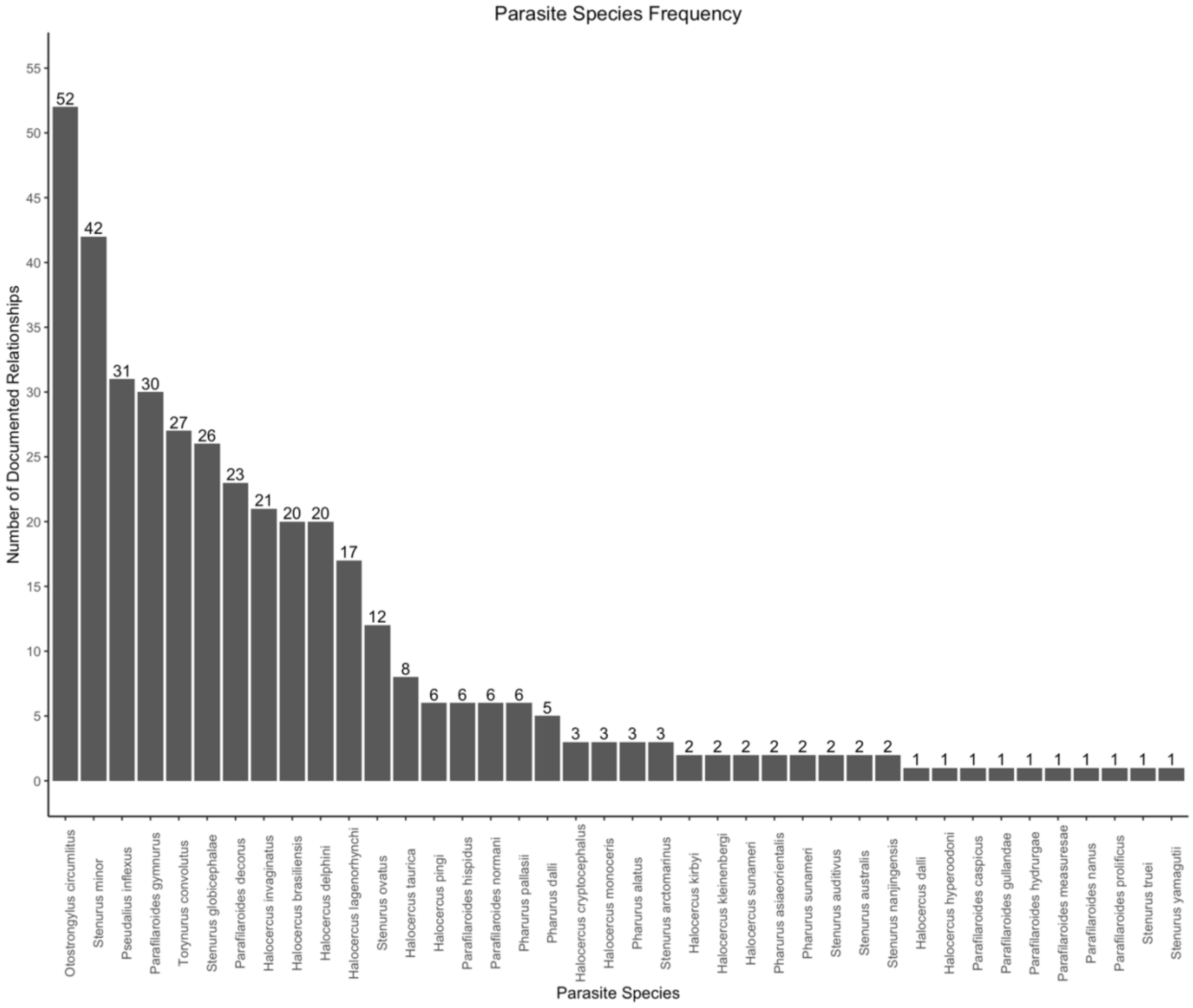


Supplementary Figure 1: Frequency of metastrongyle species documented.

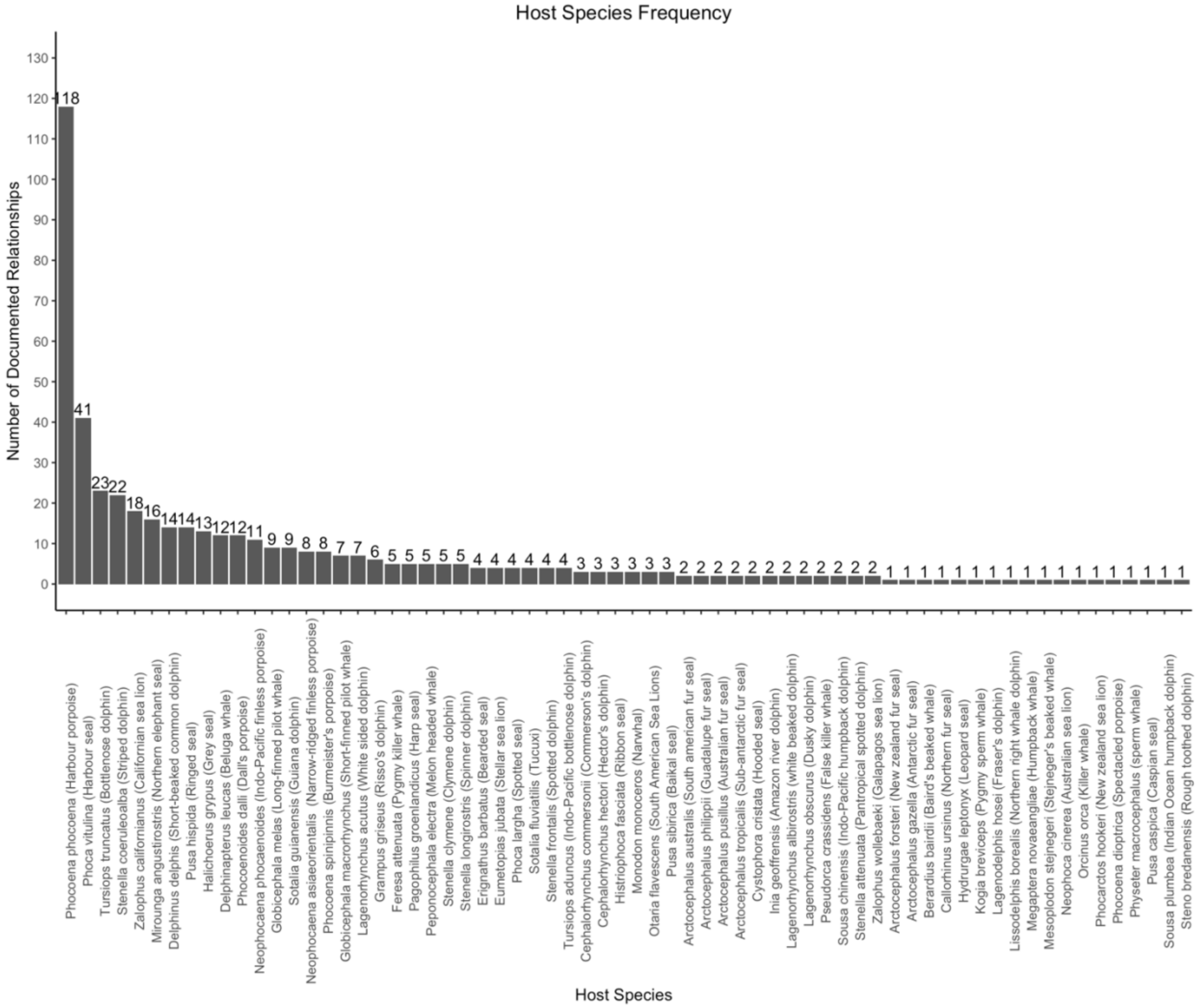


Supplementary Figure 2: Frequency of marine mammal host species documented.

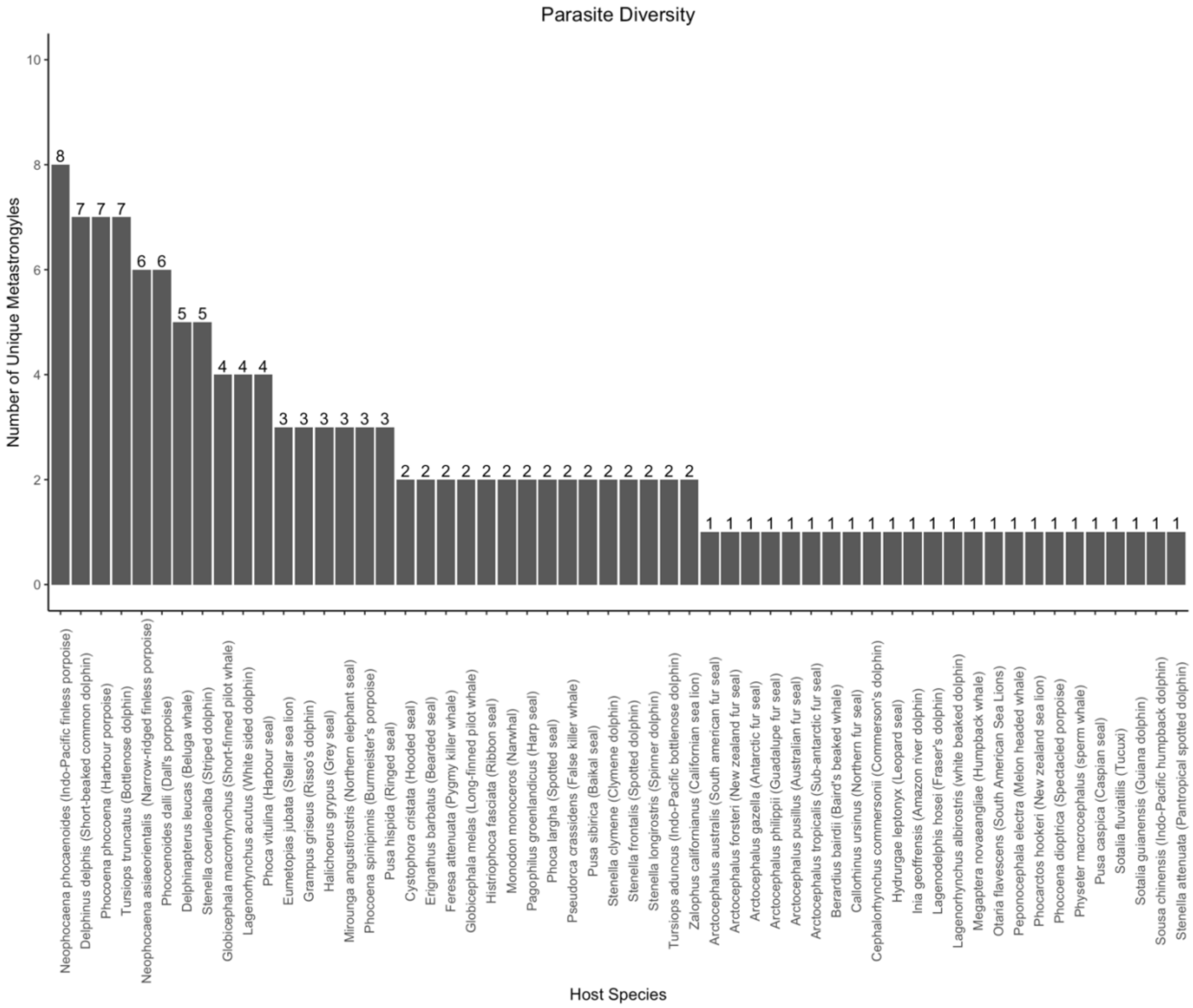


Supplementary Figure 3: Parasite diversity for metastrongyles in marine mammals (number of unique metastrongyle species hosted by each marine mammal).

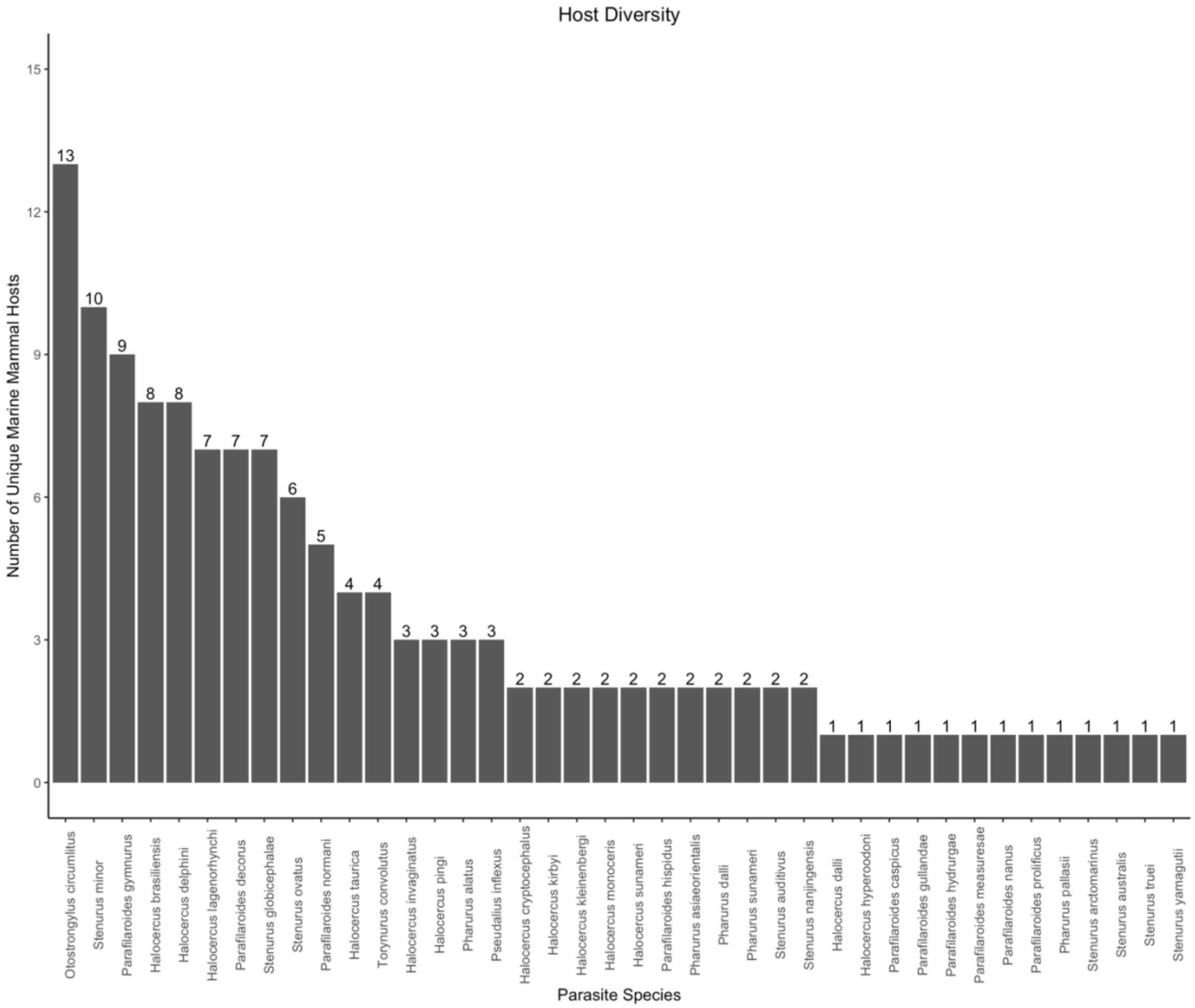


Supplementary Figure 4: Host diversity for metastrongyles in marine mammals (number of host species parasitized by each metastrongyle).
